# The oral drug nitazoxanide restricts SARS-CoV-2 infection and attenuates disease pathogenesis in Syrian hamsters

**DOI:** 10.1101/2022.02.08.479634

**Authors:** Lisa Miorin, Chad E. Mire, Shahin Ranjbar, Adam J. Hume, Jessie Huang, Nicholas A. Crossland, Kris M White, Manon Laporte, Thomas Kehrer, Viraga Haridas, Elena Moreno, Aya Nambu, Sonia Jangra, Anastasija Cupic, Marion Dejosez, Kristine A. Abo, Anna E. Tseng, Rhiannon B. Werder, Raveen Rathnasinghe, Tinaye Mutetwa, Irene Ramos, Julio Sainz de Aja, Carolina Garcia de Alba Rivas, Michael Schotsaert, Ronald B. Corley, James V. Falvo, Ana Fernandez-Sesma, Carla Kim, Jean-François Rossignol, Andrew A. Wilson, Thomas Zwaka, Darrell N. Kotton, Elke Mühlberger, Adolfo García-Sastre, Anne E. Goldfeld

**Author notes:** To whom correspondence should be addressed: Lisa Miorin; Anne E. Goldfeld. These authors contributed equally (Lisa Miorin, Chad E. Mire, Shahin Ranjbar).

## Abstract

A well-tolerated and cost-effective oral drug that blocks SARS-CoV-2 growth and dissemination would be a major advance in the global effort to reduce COVID-19 morbidity and mortality. Here, we show that the oral FDA-approved drug nitazoxanide (NTZ) significantly inhibits SARS-CoV-2 viral replication and infection in different primate and human cell models including stem cell-derived human alveolar epithelial type 2 cells. Furthermore, NTZ synergizes with remdesivir, and it broadly inhibits growth of SARS-CoV-2 variants B.1.351 (beta), P.1 (gamma), and B.1617.2 (delta) and viral syncytia formation driven by their spike proteins. Strikingly, oral NTZ treatment of Syrian hamsters significantly inhibits SARS-CoV-2-driven weight loss, inflammation, and viral dissemination and syncytia formation in the lungs. These studies show that NTZ is a novel host-directed therapeutic that broadly inhibits SARS-CoV-2 dissemination and pathogenesis in human and hamster physiological models, which supports further testing and optimization of NTZ-based therapy for SARS-CoV-2 infection alone and in combination with antiviral drugs.

## Introduction

The ongoing COVID-19 pandemic has resulted in more than 396 million cases and 5.7 million deaths worldwide (Dong et al., 2020). An easily deployable, well-tolerated, inexpensive, orally active drug that is safe in adults and children to treat or inhibit disease progression would be a significant advance in the fight against COVID-19. Nitazoxanide (NTZ) is an FDA-approved and well-tolerated oral therapy originally developed for parasitic infection that has been used for treatment of *Giardia*- and *Cryptosporidium*-associated diarrhea in millions of adults and children (Doumbo et al., 1997; Hussar, 2004; Rossignol et al., 1998). In addition, both NTZ and its circulating metabolite tizoxanide (TIZ) have been shown to inhibit a diverse array of viruses, including human coronaviruses *in vitro* (Rossignol, 2014; Wang et al., 2020).

Our previous studies showed that NTZ functions as a host-directed therapeutic (HDT) that inhibits Ebola virus (EBOV) and vesicular stomatitis virus (VSV) by amplifying host cytosolic RNA sensing and type I interferon (IFN) signaling pathways and by inducing the cell stress and antiviral phosphatase GADD34 (Jasenosky et al., 2019). These studies, which showed that NTZ creates a broad “antiviral milieu” capable of overcoming virus-specific immune evasion strategies (Jasenosky et al., 2019; Ranjbar et al., 2019), led us to test NTZ’s ability to inhibit severe acute respiratory syndrome coronavirus 2 (SARS-CoV-2) replication and pathogenesis in physiologically relevant preclinical models of infection.

Here, we show that NTZ significantly inhibits SARS-CoV-2 replication in human and primate cell lines including human alveolar epithelial type 2 cells (iAT2s) derived from induced pluripotent stem cells (iPSCs). We also show that NTZ broadly inhibits replication of SARS-CoV-2 variants of concern and SARS-CoV-2 Spike protein-induced syncytia formation. Furthermore, when administered in combination with the antiviral remdesivir (RDV), an intravenous drug that directly targets viral replication via inhibition of the RNA-dependent RNA polymerase (RdRp) (Pruijssers et al., 2020), NTZ synergistically inhibits SARS-CoV-2 growth.

Importantly, our study validates the potential of NTZ as a repurposed HDT against COVID-19 in the Syrian hamster infection model. As compared to vehicle-treated infected hamsters, oral administration of NTZ twice-daily resulted in a significant decrease in SARS-CoV-2 morbidity as characterized by both decreased weight loss and lung pathology with dramatic reduction of SARS-CoV-2 viral spike dissemination and syncytia formation. Altogether, these results provide impetus to test and optimize an NTZ-based therapy especially in combination with other promising orally bioavailable antiviral drugs that directly target viral replication.

## Results

### NTZ and TIZ inhibit SARS-CoV-2 infection in Vero E6 cells in a dose-dependent manner

To determine whether NTZ and its active metabolite TIZ inhibit SARS-CoV-2 replication, we first performed antiviral assays in Vero E6 cells, which are well known to support productive infection with SARS-CoV-2 (Matsuyama et al., 2020) (Suppl. Fig. 1A). Vero E6 cells were thus exposed to increasing concentrations of NTZ, TIZ, or vehicle (DMSO) for 4 hrs before infection with SARS-CoV-2 (isolate USA-WA1/2020) at MOI of 0.025 (see schema in Fig. 1A), and as a positive control, cells were treated in parallel with different concentrations of the RdRp inhibitor RDV. After 48 hrs, the percent of infection across the different conditions was determined by immunostaining for the viral nucleoprotein (N) using a Celigo (Nexcelcom) imaging cytometer and was used to calculate the 50% inhibitory concentration (IC_50_) of NTZ as previously described (Alwan et al., 1988). The cytotoxicity of NTZ, TIZ, and RDV was evaluated by performing MTT assays on uninfected cells treated with the same compound dilutions in parallel with the antiviral assay. We found that NTZ and TIZ possess strong antiviral activity against SARS-CoV-2 in Vero E6 cells, with IC_50_s of 4.04 μM and 3.62 μM, respectively, without exhibiting cytotoxicity (Fig. 1B).

**Fig. 1.**
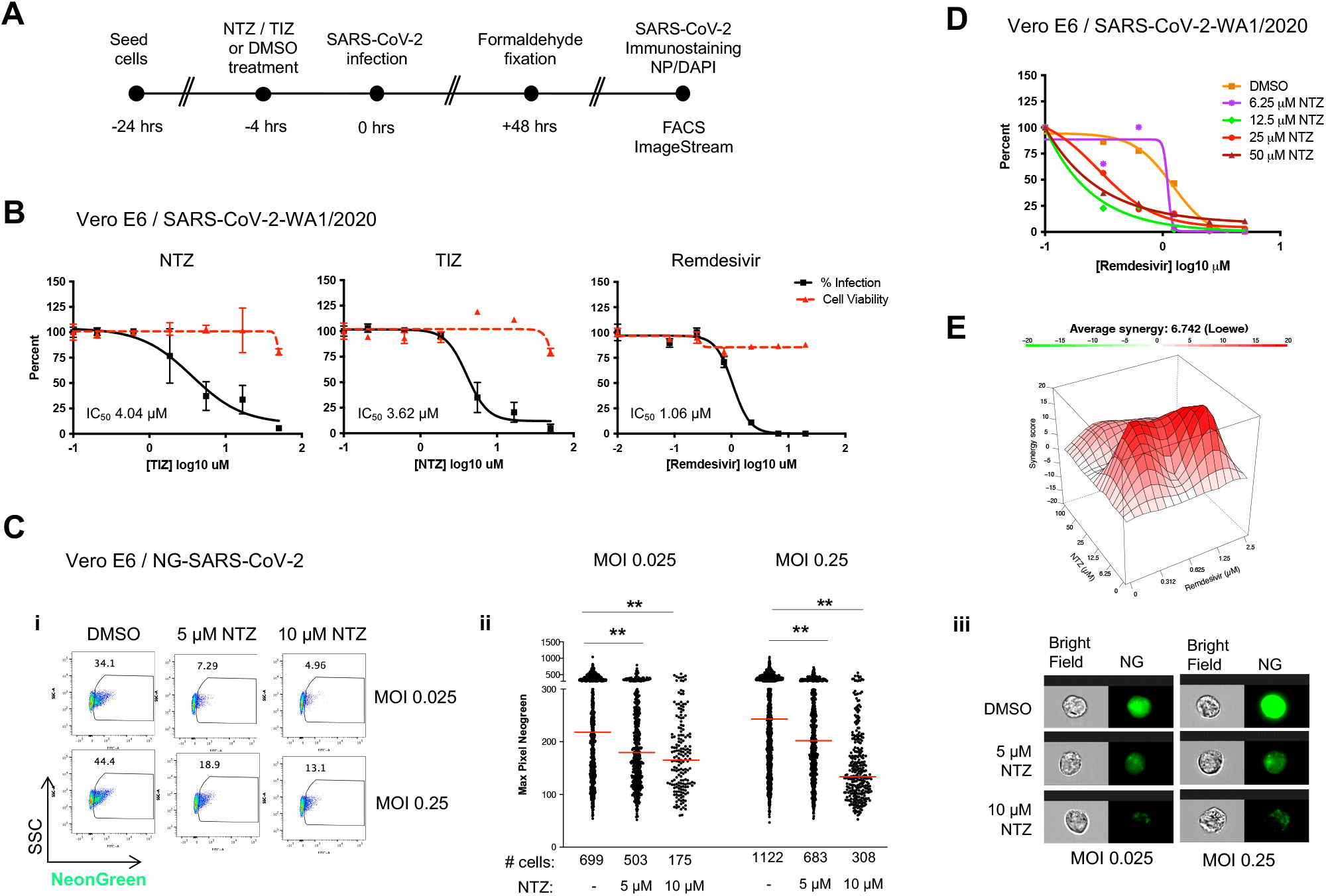
SARS-CoV-2 replication is inhibited by NTZ or TIZ in Vero E6 cells. **A**. Schema of the antiviral assay. **B**. Percent inhibition of SARS-CoV-2 replication and cytotoxicity assay in Vero E6 cells in the presence of the indicated drugs: NTZ (nitazoxanide), TIZ (tizoxanide), Remdesivir (RDV). Vero E6 cells were infected with 100 PFU (MOI 0.025) of SARS-CoV-2 (isolate USA-WA1/2020) in the presence of increasing concentrations of drug for 48 hrs, after which viral replication was measured by NP immunostaining as described in Materials and Methods. In all panels, viral infectivity is shown as a solid black line and cell toxicity as a dashed red line. Calculated IC_50_ is indicated in the bottom-left corner of each plot. RDV is included as a standard of care control. **C**. Quantitative inhibition of NeonGreen SARS-CoV-2 in Vero E6 cells measured by flow cytometry and quantitative imaging flow cytometry using the ImageStream platform. Vero E6 cells were pretreated with 5 or 10 μM NTZ or carrier (DMSO) for 4 hrs and then infected with NeonGreen (NG)-SARS-CoV2 at an MOI of 0.025 or 0.25. At 48 hrs post-infection cultures were fixed for 24 hrs before FACS and ImageStream analysis. (i) The percentage of infection as measured by FACS. Experiments were performed at least three independent times. (ii) Dot plot displaying the quantification of NG pixels as a representation of NG-SARS-CoV-2 fluorescence in infected cells (among the 3000 cells acquired). Asterisks indicate significant differences in NTZ-treated samples as compared to DMSO control by two-tailed Mann-Whitney test. \*\**P*<0.001 for each comparison. The number of infected cells in each experimental condition is shown at the bottom of the panel. We found significant reduction of infected cells in NTZ-treated cultures at both MOIs and both concentrations of NTZ (\*\*\**P*<0.0001 for each comparison) by Fisher exact test using Stata 12 software. Experiments were performed at least three independent times. (iii) Images of representative NeonGreen-SARS-CoV-2 infected cells were selected from the median levels of pixels shown in the analysis in panel (ii). **D**. Antiviral activity curves by RDV in the presence of increasing concentrations of NTZ. **E**. Synergy landscapes and combination scores generated by the Loewe method using SynergyFinder software.

Using a NeonGreen (NG) expressing SARS-CoV-2 (USA-WA1/2020) recombinant virus (NG-SARS-CoV-2) (Xie et al., 2020), the effect of NTZ on SARS-CoV-2 replication was next assessed by FACS and by imaging flow cytometry using the ImageStream platform (Doan et al., 2018; Haridas et al., 2017; Ranjbar et al., 2019). Cells were pre-treated with 5 or 10 μM NTZ for 4 hrs and then infected with NG-SARS-CoV-2 at two different MOIs (0.025 and 0.25) for 48 hrs. In agreement with our findings using the USA-WA1/2020 strain in the antiviral assays above (Fig. 1B), we observed dose-dependent NTZ inhibition of the percentage of NG-SARS-CoV-2-infected cells at both MOIs by FACS analysis (Fig. 1C-i).

Imaging flow cytometry allows for high-throughput evaluation of SARS-CoV-2 infection at the single cell level (Doan et al., 2018; Haridas et al., 2017; Ranjbar et al., 2019) and quantification of intracellular fluorescence produced by NG-labeled SARS-CoV-2. Using the ImageStream platform, we thus next acquired approximately 3000 cells from vehicle- and NTZ-treated NG-SARS-CoV-2-infected cultures and measured NG fluorescence in each experimental condition. NTZ pre-treatment for 4 hrs prior to SARS-CoV-2 infection resulted in significant inhibition of NG expression at both concentrations of NTZ (5 or 10 μM) and at both MOIs (0.025 and 0.25) (p<0.001 for each comparison) after 48 hrs (Fig. 1C-ii). We note that representative images of cells that had pixel values at the median level of each condition (marked by the red bar) are displayed showing the NTZ-dependent decreased fluorescence in the infected cells (Fig. 1C-iii). Strikingly, we also observed a significant decrease of the number of infected cells at both NTZ concentrations and both MOIs tested (Fig. 1C-ii) (p<0.0001) for each comparison.

Given the robust antiviral activity of NTZ in Vero E6 cells, and based on their different mechanisms of action, we next assessed NTZ’s ability to synergize with RDV, which was a standard of care drug treatment to shorten time of COVID-19 recovery (Beigel et al., 2020). To test this hypothesis, we performed combination assays in Vero E6 cells and found that NTZ significantly increases the ability of RDV to inhibit viral infection (Fig. 1D). The results of the combination assay were then further analyzed using SynergyFinder to generate a synergy landscape and combination score by the Loewe model (Fig. 1E) (Ianevski et al., 2020). In this model, a synergistic interaction between drugs is indicated by a score greater than +10, while an additive interaction has a score between -10 and +10, and an antagonistic interaction has a score lower than -10. Notably, the landscape for the interaction of RDV with NTZ shows synergy scores greater than +10 at low NTZ concentrations (6.26 μM) (Fig. 1E), denoting moderate synergy of the two drugs in the inhibition of SARS-CoV-2 growth. These data are consistent with a study showing that NTZ synergistically enhances the ability of RDV to reduce cytopathic effect (CPE) (Bobrowski et al., 2021). All together, these data show that in Vero E6 cells NTZ robustly inhibits SARS-CoV-2 replication and forms an inhibitory synergistic compound pair with RDV.

### Validation of NTZ antiviral activity in human cell lines

We next evaluated the ability of NTZ to inhibit SARS-CoV-2 replication in the human cell line A549 transduced with angiotensin-converting enzyme 2 (Ace2). After confirming that the Ace2-A549 cells support SARS-CoV-2 replication (Suppl. Fig. 1B), we pre-treated these cells with increasing concentrations of NTZ, TIZ, RDV, or vehicle (DMSO) for 4 hrs before infection with SARS-CoV-2 (USA-WA1/2020) at an MOI of 0.1. At 48 hrs post-infection, the IC_50_ and cytotoxicity of each compound was determined as described above. We observed strong antiviral activity of NTZ (IC_50_ 1.695 μM) and TIZ (IC_50_ 1.322 μM) against SARS-CoV-2 infection in Ace2-A549 with no cytotoxicity (Fig. 2A). In addition, we also evaluated the effect of NTZ on SARS-CoV-2 replication in another human cell line, Ace2-expressing HEK293T cells (Ace2-HEK293T) (Miorin et al., 2020). NTZ similarly inhibited SARS-CoV-2 replication in a dose-dependent manner in Ace2-HEK293T with an IC_50_ of 2.2 μM (Suppl. Fig. 2).

**Fig. 2.**
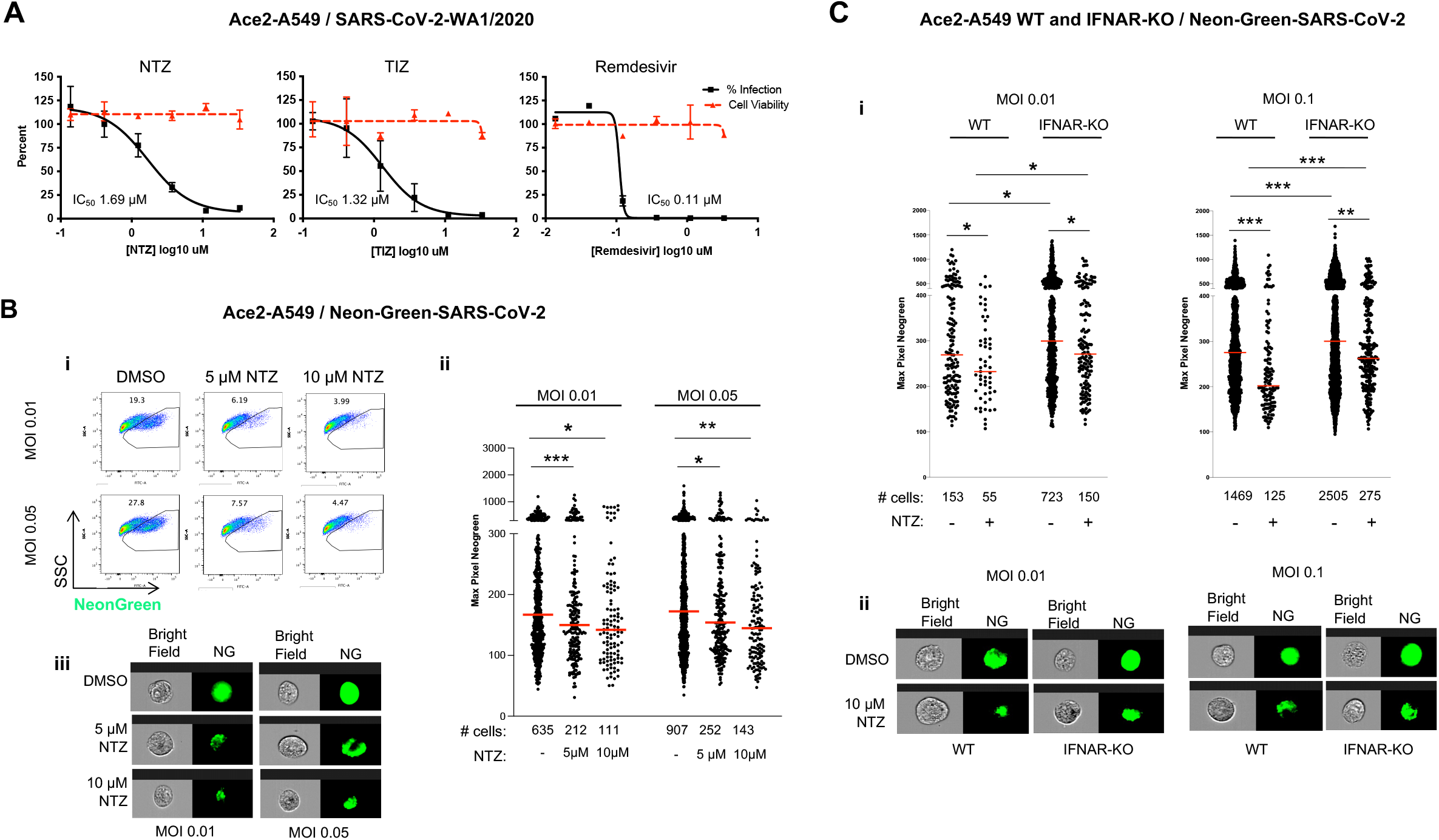
Antiviral activity of NTZ and TIZ in Ace2-A549 cells. **A**. Percent inhibition of SARS-CoV-2 replication and cytotoxicity assay in Ace2-A549 cells in the presence of NTZ, TIZ, or RDV. Cells were treated with the indicated drug for 4 hrs prior to infection with SARS-CoV-2 (isolate USA-WA1/2020) at MOI 0.1 for 48 hrs. Viral replication was measured by NP immunostaining as described in Materials and Methods. In all panels, viral infectivity is shown as a solid black line and cell toxicity as a dashed red line. Calculated IC_50_ is indicated at the bottom of each plot. RDV is included as a standard of care control. **B**. Quantitative inhibition of NG-SARS-CoV-2 in Ace2-A549 cells measured by flow cytometry and quantitative imaging flow cytometry using the ImageStream platform. Ace2-A549 cells were pretreated with 5 or 10 μM NTZ or carrier (DMSO) for 4 hrs before infection at an MOI of 0.05 or 0.01. At 48 hrs post-infection cells were fixed and evaluated by FACS and by quantitative imaging flow cytometry as described in the legend to Fig. 1. (i) The percentage of infection as measured by FACS. Experiments were performed at least three independent times. (ii) Dot plot displaying the quantification of NG pixels as a representation of NG-SARS-CoV-2 fluorescence in infected cells (among the 3000 cells acquired). Asterisks indicate significant differences as compared to DMSO control by two-tailed Mann-Whitney test: \*\**P*<0.005 and \**P*<0.05. The number of infected cells in each experimental condition is shown at the bottom of the panel. We found significant reduction of infected cells in NTZ-treated cultures at both MOIs and both concentrations of NTZ (\*\*\**P*<0.0001 for each comparison). Experiments were performed at least three independent times. (iii) Images of representative NeonGreen-SARS-CoV-2 infected cells were selected from the median levels of pixels shown in the analysis in panel (ii). **C**. NTZ’s inhibitory effect on NG-SARS-CoV-2 growth in wild-type (WT) and IFNAR-KO Ace2-A549 cells. (i) Plots displaying the quantification of NG pixels as in (B). The number of infected cells in each experimental condition is shown at the bottom of the panel. We found significant reduction of infected cells in NTZ treated cultures at both MOIs and both concentrations of NTZ (\*\*\**P*<0.0001 for each comparison) by Fisher exact test using Stata 12 software. Experiments were performed at least three independent times. (ii) Images of representative NeonGreen-SARS-CoV-2 infected cells were selected from the median levels of pixels shown in the analysis in panel. In all panels, asterisks indicate significant differences as compared to DMSO control by two-tailed Mann-Whitney test: \**P*<0.05, \*\**P*<0.005, \*\*\**P*<0.0001.

We next pretreated Ace2-A549 cells with 5 or 10 μM NTZ for 4 hrs followed by infection with NG-SARS-CoV-2 (MOI 0.01 and 0.05) for 48 hrs. Analysis by FACS showed that NTZ reduced the percentage of NG-SARS-CoV-2-infected Ace2-A549 cells at both NTZ concentrations and both MOIs (Fig. 2B-i). Furthermore, 5 μM NTZ significantly reduced both the median levels of NG-SARS-CoV2 fluorescence in infected cells and the number of infected cells (Fig. 2B-ii). Images of individual Ace2-A549 cells with pixel numbers that fell at the median level of pixels for the population are displayed in Figure 2B-iii.

Interestingly, the IC_50_ for NTZ in both human cell lines (Ace2-A549: 1.695 μM; Ace2-HEK293T: 2.2 μM) were lower than the IC_50_ in Vero E6 cells (4.444 μM). Notably, Vero cells are deficient in type I IFN production (Emeny and Morgan, 1979) due to a deletion on chromosome 12 resulting in loss of the type I IFN gene cluster (Osada et al., 2014). To interrogate whether signaling through the type I IFN receptor (IFNAR) plays a role in NTZ’s ability to inhibit SARS-CoV-2 infection in human cells, we compared NTZ antiviral activity in wild-type and IFNAR knock-out (KO) Ace2-A549 cells. To verify that IFNAR signaling was abrogated in the knock-out cells, we showed that the IFN-dependent phosphorylation of STAT1 and STAT2 (Suppl. Fig. 3A) and transcription of the interferon stimulated gene (ISG) IFITM3 (Suppl. Fig. 3B) were both abolished in the IFNAR-KO Ace2-A549 cells as expected. In addition, we confirmed that Ace2 expression was equivalent in the two cell lines (Suppl. Fig. 3A).

**Fig 3.**
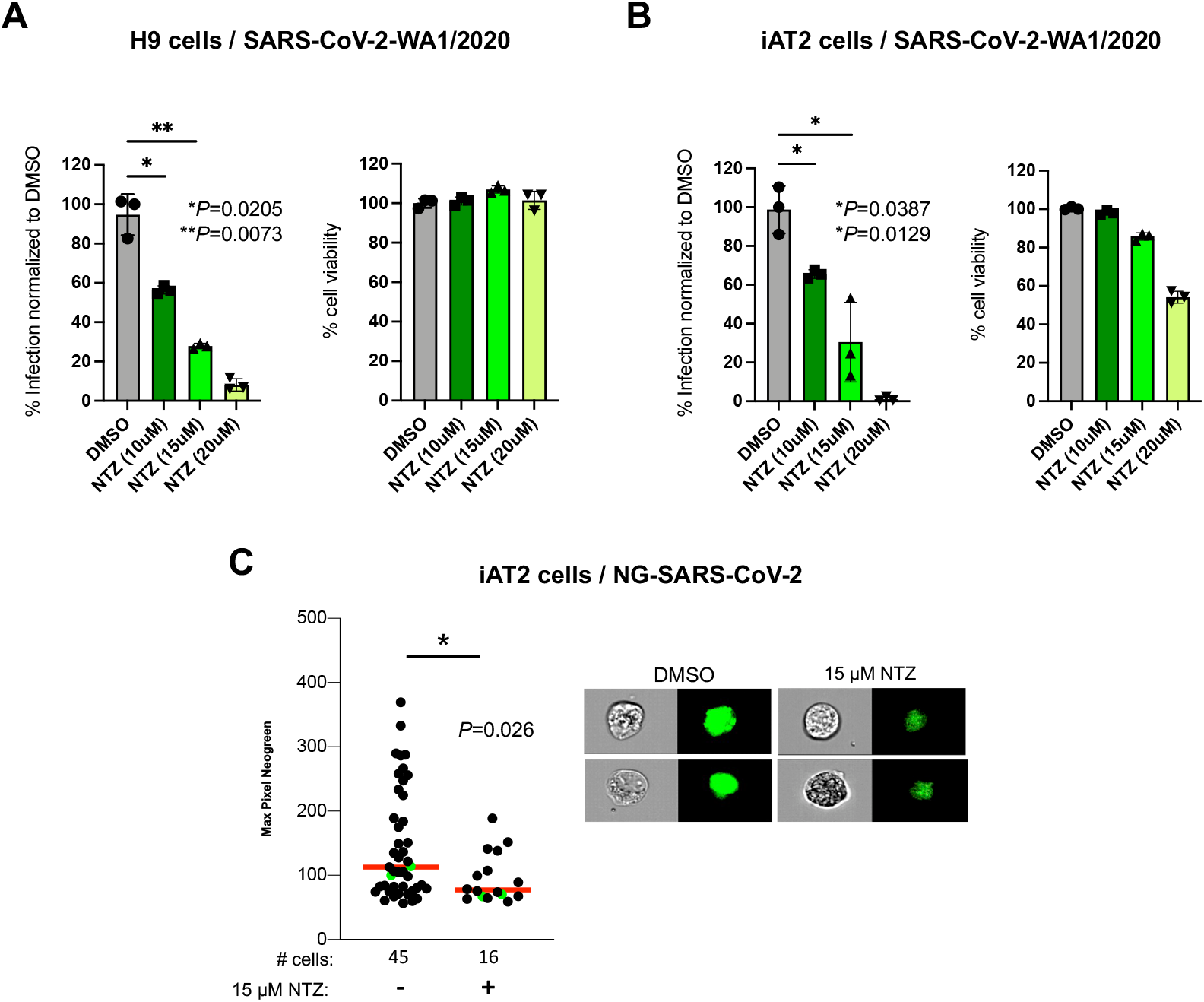
NTZ is highly active against SARS-CoV-2 infection in stem cell-derived human lung alveolar epithelial cultures. **A**. H9-derived pneumocyte-like cells were infected with SARS-CoV-2 (isolate USA-WA1/2020) at an MOI of 0.1 in the presence of the indicated concentrations of NTZ or vehicle (DMSO). All cells were pretreated for 4 hrs and NTZ or DMSO were maintained in the media throughout the experiment. SARS-CoV-2 infection (NP immunostaining) and cell viability (MTT assay) were measured at 48 hrs. Asterisks indicate statistically significant differences as shown on the graph. **B**. iPSC-derived alveolar epithelial type 2 cells (iAT2s) were infected with SARS-CoV-2 (isolate USA-WA1/2020) at an MOI of 0.5 in the presence of the indicated concentrations of NTZ or vehicle (DMSO) as described in Materials and Methods. Percent infection was measured by NP immunostaining and cell viability by trypan blue exclusion performed on infected and treated cells. Asterisks indicate statistically significant differences as shown on the graph. Experiments were performed at least three independent times. **C**. Quantitative imaging flow cytometry IDEAS analysis of NG-SARS-CoV-2-infected iAT2 in the presence of vehicle (DMSO) or NTZ (15 μM). 3000 cells were acquired on an ImageStream^X^ Mark II instrument and evaluated for pixel intensity of NeonGreen (left). The median cellular fluorescence of the NG-SARS-CoV-2-infected iAT2 cells was significantly reduced in the NTZ-treated vs. mock-treated cultures (p=0.026). The reduction of the number of infected cells in NTZ-treated cultures was significant (\*\*\**P*<0.0001) by Fisher exact test using Stata 12 software. Representative images of intracellular NG levels at the median range marked by the red line in the dot plot (right).

The wild-type and IFNAR-KO Ace2-A549 cells were then treated with 10 μM NTZ or with DMSO for 4 hrs followed by infection with NG-SARS-CoV-2 at an MOI of 0.1 or 0.01 for 48 hrs. Notably, IFNAR-KO Ace2-A549 cells showed a significant increase in the median level of NG fluorescence and also in the numbers of SARS-CoV-2-infected cells at both NTZ concentrations and MOIs tested (Fig. 2C-i). Thus, IFNAR signaling restricts SARS-CoV-2 growth and infection in Ace2-A549 cells. Interestingly, NTZ significantly inhibited both the level of intracellular NG-SARS-CoV-2 fluorescence and the number of infected cells in both WT and IFNAR-KO Ace2-A549 cells, however its ability to inhibit SARS-CoV-2 replication and the number of infected cells was reduced in the IFNAR-KO cells (Fig. 2C-i). Indeed, the median NG fluorescence and the number of infected cells remained significantly higher in the NTZ-treated IFNAR-KO cells compared to the NTZ-treated parental Ace2-A549 cells at both MOIs. Taken together, these results show: (i) IFNAR signaling restricts SARS-CoV-2 replication; (ii) NTZ can inhibit SARS-CoV-2 in the absence of IFNAR signaling; and (iii) IFNAR signaling is required for optimal NTZ inhibition of SARS-CoV-2 growth and infection.

### NTZ inhibits SARS-CoV-2 replication in physiologic human pluripotent stem cell-derived cell models

SARS-CoV-2 targets and infects ciliated airway epithelial cells and type 2 pneumocytes in alveolar regions of the lung (Hou et al., 2020). To evaluate the ability of NTZ to restrict SARS-CoV-2 replication in a physiologically relevant cell type, we used two independently derived human pneumocyte *in vitro* models that are generated by the directed differentiation of human pluripotent stem cells (PSCs), the H9 embryonic stem cell line and SPC2-ST-B2 induced pluripotent stem cells (iPSCs) (Hurley et al., 2020). Both H9-derived pneumocyte-like cells (Riva et al., 2020) and the iPSC-derived alveolar epithelial type 2 cells (iAT2s) (Huang et al., 2020) have previously been shown to support SARS-CoV-2 replication. Strikingly, pre-treatment of H9-pneumocytes (Fig. 3A) or iAT2 cells (Fig. 3B) with NTZ for 4 hrs prior to infection with SARS-CoV-2 (isolate USA-WA1/2020) resulted in a significant dose-dependent inhibition of virus replication without significant cytotoxicity. In addition, NTZ’s antiviral activity against SARS-CoV-2 in iAT2 cells was also validated using the NG-SARS-CoV-2 recombinant virus and the ImageStream platform. We found that NTZ significantly inhibited NG-SARS-CoV-2 fluorescence (p=0.026) and the number of NG-SARS-CoV-2 infected cells in the culture (p<0.0001) (Fig. 3C). We note that the PSCs from which H9 and iAT2 cells are derived are from two unrelated individuals. Thus, our findings are not specific to an individual and indicate that NTZ inhibits both SARS-CoV-2 replication and *de novo* infection of bystander cells in human alveolar epithelial cell cultures that reflect physiological infection.

### NTZ inhibits virus replication of SARS-CoV-2 variants of concern

Different SARS-CoV-2 variants have contributed to successive waves of the COVID-19 epidemic due to their increased transmissibility and virulence, as well as the waning of vaccine-mediated immunity (Harvey et al., 2021). To determine NTZ’s activity against three of the recently emerged variants of concern (https://www.who.int/en/activities/tracking-SARS-CoV-2-variants/), we treated Vero E6 cells with NTZ for 4 hrs and then infected them with SARS-CoV-2 beta (B.1.351), gamma (P.1), or delta (B.1617.2). As a control, cells were also infected with the original SARS-CoV-2 WA1 strain used in our studies described above. NTZ strongly inhibited the replication of each variant tested similar to its ability to inhibit SARS-CoV-2 WA1 (Fig. 4A). Thus, NTZ is able to inhibit different viral variants with divergent spike proteins that are associated with increased virulence and transmissibility.

**Fig 4.**
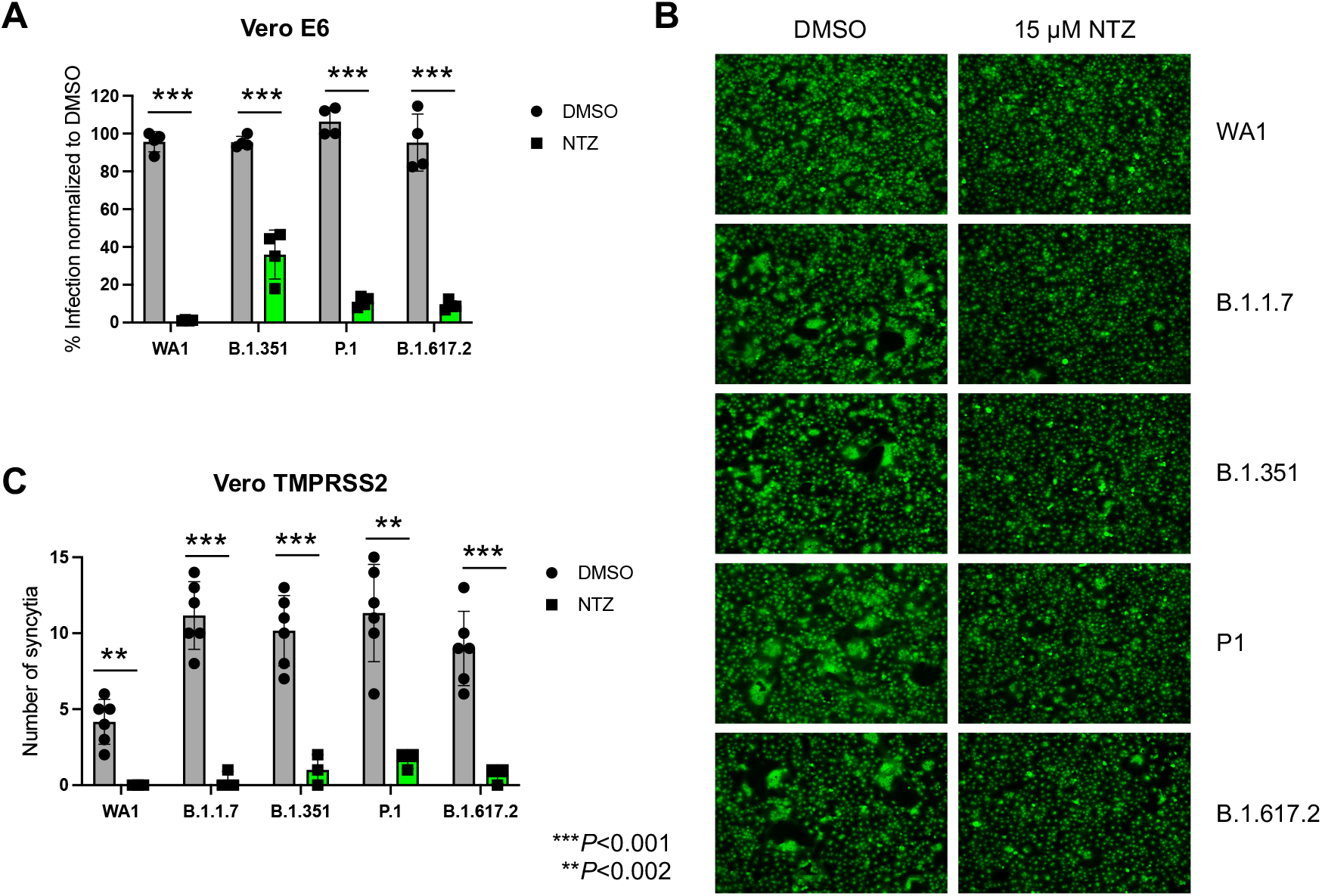
NTZ inhibits replication of different SARS-CoV-2 VOCs and strongly reduces spike-mediated syncytia formation. **A**. Vero E6 cells were treated with DMSO or 15 μM NTZ four hrs before infection with the indicated SARS-CoV-2 isolates at an MOI of 0.025 for 48 hrs. Viral replication was measured by NP immunostaining and quantified using a Celigo imaging cytometer as described in Materials and Methods. **B**. Vero TMPRSS2 cells were reverse transfected with 100 ng of pCAGGS plasmids expressing the spike protein from the SARS-CoV-2 variants of concern indicated in the figure. Four hours after transfection, medium containing transfection complexes was removed and replaced by medium containing 2% FBS and either DMSO or 15 μM NTZ. After 24 hrs, cells were fixed and stained with HCS Cell Mask stain (Life Technologies). **C**. Quantification of spike-induced syncytia formation from (B). Syncytia were counted microscopically in triplicate wells. In all panels, asterisks indicate statistically significant differences (\*\*\**P*<0.001).

Intriguingly, high-throughput screening of over 3,000 FDA/EMA-approved drugs recently identified niclosamide, which is structurally similar to NTZ (Stachulski et al., 2021), as an effective inhibitor of spike-induced syncytia formation (Braga et al., 2021). To test the hypothesis that NTZ restricts SARS-CoV-2 dissemination by directly inhibiting spike-mediated fusion, we overexpressed spike protein from SARS-CoV-2 (USA-WA1/2020) and SARS-CoV-2 variants alpha (B.1.1.7), beta (B.1.351), gamma (P.1), and delta (B.1617.2) in Vero-TMPRSS2 cells. Four hours after transfection, 15 μM NTZ or vehicle (DMSO) was added to the cultures and 24 hrs later cells were fixed to evaluate syncytia formation. NTZ broadly and significantly inhibited syncytia formation induced by overexpression of spike from the USA-WA1/2020 strain and from each variant tested in this assay. Consistent with their increased virulence compared to the USA-WA1/2020 strain, the four variants exhibited significantly higher levels of syncytia formation (Fig. 4B), concordant with a recent report (Escalera et al., 2021). These data indicate that NTZ targets a cellular process mediating spike-induced cell-cell fusion that is not significantly affected by the spike mutations in the variants tested.

### Oral administration of NTZ significantly reduces SARS-CoV-2 replication and disease pathogenesis in Syrian hamsters

Given the promising antiviral activity of NTZ *in vitro*, we next evaluated its ability to inhibit SARS-CoV-2 infection and disease *in vivo*. We chose the Syrian hamster model of SARS-CoV-2 infection, where animals develop severe pneumonia and clinical outcomes (i.e., peak weight loss around 5 days and complete resolution by 14 days post infection), which are predictable (Chan et al., 2020; Imai et al., 2020; Sia et al., 2020). We used a new extended-release formulation of NTZ (NT-300) currently under phase III evaluation (Rossignol et al., 2021), which reaches peak levels 6-8 hrs after ingestion (Haffizulla et al., 2014).

Hamsters were split into 3 groups (12 animals/group): (i) naïve (no infection and no treatment); (ii) infected with SARS-CoV-2 (300pfu) with vehicle (PBS) treatment delivered twice daily (BID) to control for hamster manipulation during oral gavage; (iii) infected with SARS-CoV-2 (300 pfu) and with NTZ (300 mg/kg/day) delivered BID by oral gavage. The first NTZ or vehicle dose was administered 6 hours prior to SARS-CoV-2 infection. The second dose was given 6 hours after infection and then NTZ or vehicle was dosed every 12 hours to complete 5 days of therapy. Daily weights were recorded in the 3 groups from the day of infection up until 14 days post infection (dpi). We found significant protection (p=0.025) from weight loss at 5 dpi in the NTZ-treated animals compared to animals who were treated with PBS and observed strong trends of protection at 4 dpi (p=0.056) and 6 dpi (p=0.80) (Fig. 5A). In addition, animals from each group were sacrificed at 2, 5, and 14 dpi and lungs were collected and processed for viral titer determination and histopathological evaluation. NTZ-treated SARS-CoV-2-infected animals displayed lower viral titers in lung biopsies as compared to mock treated SARS-CoV-2-infected animals (p=0.057) at 2 dpi (Fig. 5B).

**Fig 5.**
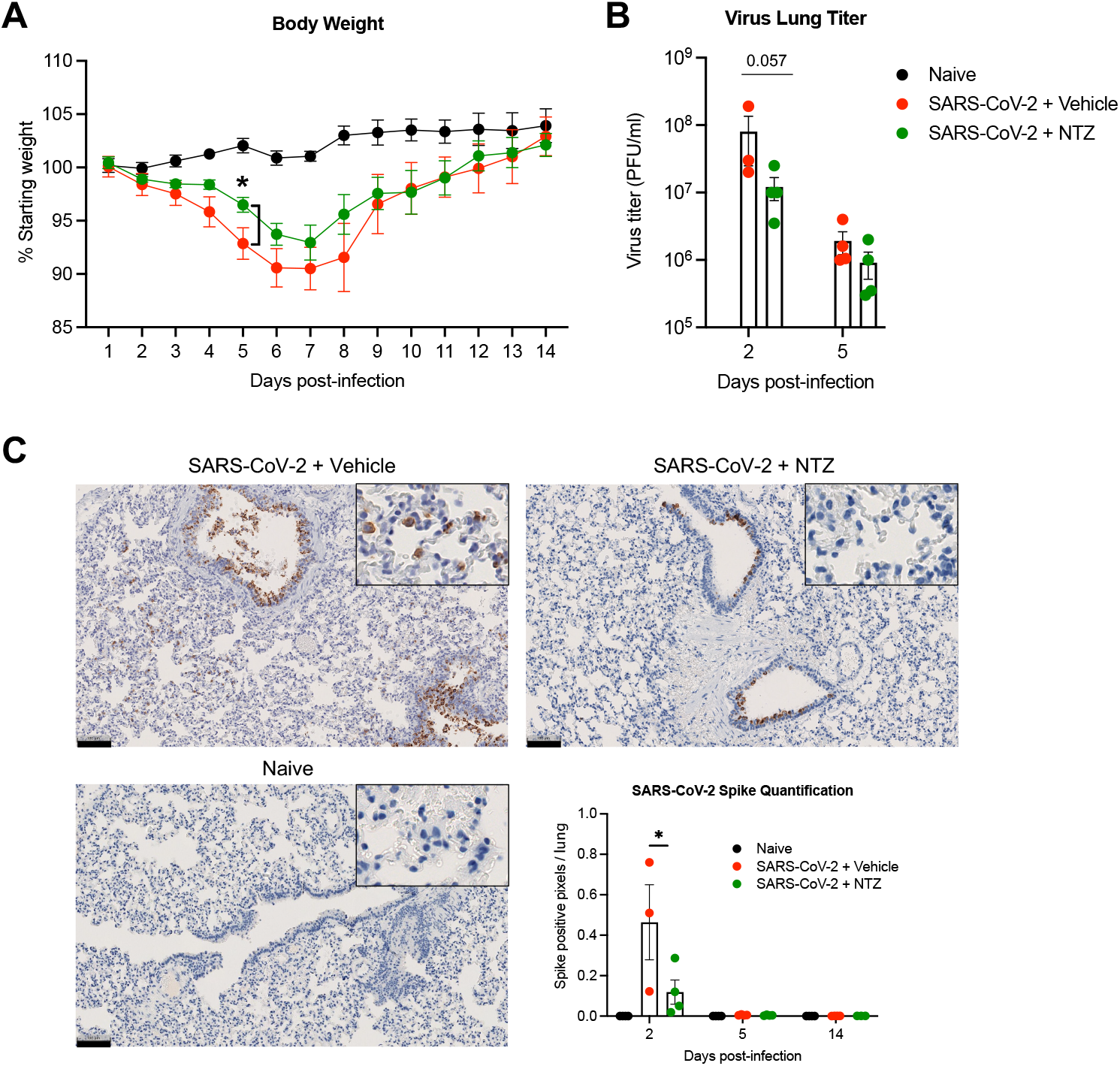
Oral delivery of NTZ significantly reduces weight loss and decreases lung viral load in the Syrian hamster model. **A**. Percent of starting body weight from the day of challenge (day 0) to 14 days post-infection in naïve and SARS-CoV-2 (isolate USA-WA1/2020)-infected Syrian hamsters treated with NTZ 300 mg/kg bi-daily or treated with vehicle by oral gavage. Four animals were euthanized on days 2, 5, and 14 for lung removal and serum collection. Day 14 was the study endpoint. Asterisk indicates statistically significant differences by one-tailed unpaired Student’s t-test. **B**. SARS-CoV-2 titer in lungs from vehicle- and NTZ-treated animals. The *P* value indicated shows the differences at day 2 by one-tailed unpaired Mann-Whitney test. Lungs from 4 animals were included in the IHC analysis for each timepoint except for the 2 dpi timepoint, where there were 3 SARS-CoV-2 PBS/vehicle-treated animals. **C**. Representative images of lung tissue collected at 2 dpi and subjected to immunohistochemistry with SARS-CoV-2 Spike-specific antibody. Insets show higher magnification of the lung interstitium, where viral antigen was only observed in the SARS-CoV-2 vehicle-treated animals. Scale bars, 100 μm. Graph shows quantification of pixels of SARS-CoV-2 Spike protein/micron of Syrian hamster lung tissue. Asterisk indicates statistically significant differences by one-tailed unpaired Student’s t-test at day 2.

In a complementary approach, we used immunohistochemistry targeting the SARS-CoV-2 Spike protein to visualize and measure the dissemination of infection within the lung. In addition, we quantified the levels of Spike protein expression in the lung by calculating the positive pixels of SARS-CoV-2 Spike immunoreactivity per micron of tissue (Fig. 5C and Suppl. Table 2). As shown in the graph in Fig. 5C, SARS-CoV-2 Spike protein expression was only detected at 2 dpi, irrespective of the cohort, and was significantly less in the NTZ-treated SARS-CoV-2-infected group (p=0.0496) as compared to the vehicle-treated SARS-CoV-2-infected cohort. This is consistent with the decline in infectious virus particles we observed at 2 dpi in the NTZ-treated animals (Fig. 5B).

While SARS-CoV-2 Spike was mostly located in bronchiole epithelium in both the vehicle- and NTZ-treated infected hamsters at 2 dpi, in the SARS-CoV-2-vehicle treatment group, Spike was more plentiful, with near-complete circumferential involvement of bronchioles that routinely disseminated to smaller distal airways with sporadic involvement of cuboidal pneumocytes interpreted to represent alveolar type 2 pneumocytes, and Spike was evident in the interstitial compartment of the lungs. By contrast, in the NTZ-treated group, almost all the SARS-CoV-2 Spike was located in the proximal airways with a more localized and segmental distribution, and it was not detected in the interstitial compartment in lungs from NTZ-treated animals (see magnification of images in Fig. 5C). By 5 dpi, there was minimal residual SARS-CoV-2 Spike observed in both the NTZ- and vehicle-treated SARS-CoV-2-infected groups, which was almost exclusively restricted to sporadic individual bronchiole epithelial cells. No SARS-CoV-2 Spike protein was detected in any cohorts by 14 dpi (data not shown).

### NTZ improves SARS-CoV-2-driven lung pathology in Syrian hamsters

Lung histopathology was also examined across the 3 experimental cohorts. Individual lung ordinal scores from 1-3 were determined for all animals to quantify severity of infection (Fig. 6B and Suppl. Table 2), and representative lung H&E images are shown (Fig. 6A and Suppl. Fig. 5). At 2 dpi the interstitial score was significantly decreased (p=0.026) in the NTZ-treated animals compared to the SARS-CoV-2-infected vehicle-treated animals (Fig. 6B). This was reflected by the absence of, or minimal, lymphohistiocytic and neutrophilic alveolar and/or septal infiltrate in the group of NTZ-treated animals (Fig. 6A). While at 2 dpi the airway score for NTZ-vs. vehicle-treated animals did not reach significance (Fig. 6C), three of the four NTZ-treated animals exhibited absence of discernible bronchiole epithelial pathology like that observed in naïve animals, with a dramatic decline in neutrophilic and histiocytic luminal infiltrate compared to vehicle-treated animals in the one animal that did display airway injury (Fig. 6A). Histologically, this was reflected by less severe and more segmental necrotizing bronchiolitis (see magnified image in Suppl. Fig. 5). Blood vessel scores for NTZ-vs. vehicle-treated animals at 2 dpi also did not reach significance, however again we note that three of the four NTZ-treated animals exhibited an absence of any detectable perivascular infiltrate similar to naïve animals.

**Fig 6.**
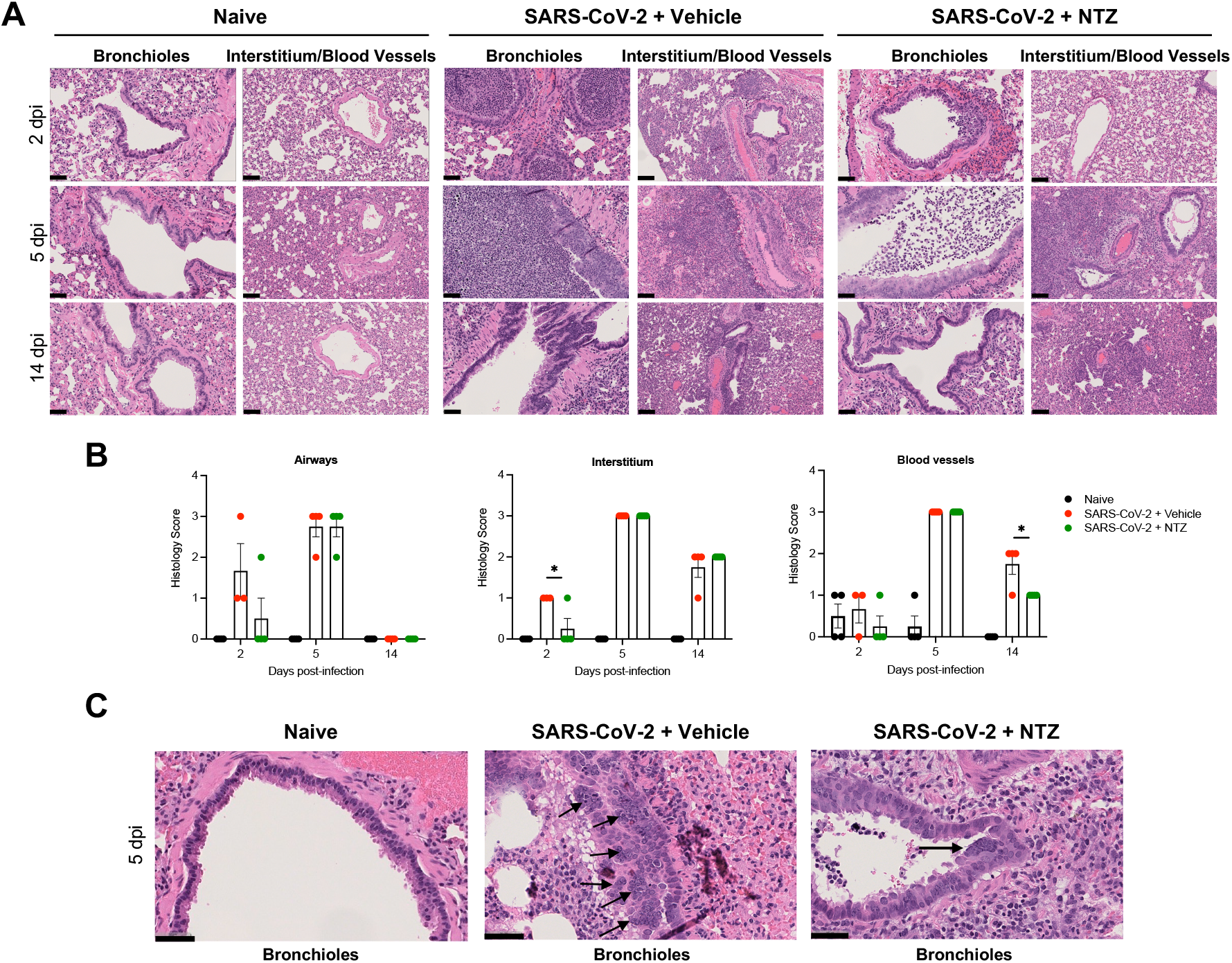
NTZ significantly prevents lung pathology SARS-CoV-2 infected Syrian hamsters. **A**. Comparison of normal lung (naïve) to pathologic changes induced by SARS-CoV-2 in vehicle-vs. NTZ-treated animals at day 2, 5, and 14 post-infection. Lung pathology at 5 dpi was severe irrespective of treatment group. **B**. NTZ significantly reduced interstitial inflammatory and perivascular scores at 2 and 14 days post-infection. Qualitatively there was decreased bronchiole epithelial injury and associated luminal inflammatory exudate. Lung pathology at 5 dpi was severe irrespective of treatment group. Asterisk indicates statistically significant differences by one-tailed unpaired Student’s t-test at day 2 for interstitium and on day 14 for blood vessels. Lungs from 4 animals were included in the scoring analysis for each timepoint except for 2 dpi, where there were 3 SARS-CoV-2 PBS/vehicle-treated animals. **C**. Bronchiole syncytial cells were observed exclusively at 5 dpi in SARS-CoV-2-inoculated animals and were more frequently observed in the vehicle treated group. Scale bars: A, 100 μm; C, 50 μm.

NTZ therapy was halted at 4 dpi with 5 days of total treatment. At 5 dpi, lung pathology scores were ubiquitously severe, with no clear distinction between the cohorts in any of the anatomical compartments examined (Figs. 6A and 6B). The precise mechanism for the uniform severity observed across the SARS-CoV-2-infected animals +/- NTZ at 5 dpi is unknown. We note that this observation mirrors findings described in an intramuscular DNA vaccine platform encoding SARS-CoV-2 Spike protein, which showed immunologic correlates of protection and a decrease in viral load in Syrian hamsters, but no clear mitigation in lung pathology at 4 dpi compared to positive controls (Leventhal et al., 2021).

At 14 dpi, no residual pathology was evident in the airways and the severity of interstitial pathology was minimal to mild in all cohorts (Fig. 6A), which was reflected by small residual clusters of AT2 hyperplasia with minimal histiocytic and neutrophilic infiltrate (Suppl. Fig. 5) with no significant difference in interstitial scores between the groups (Fig. 6B). Notably, blood vessel scores at 14 dpi were significantly decreased (p=0.012) in the NTZ-treated group compared to the vehicle-treated group (Fig. 6B), where all four NTZ-treated animals had a clear decrease in residual perivascular lymphohistiocytic infiltrates at 14 dpi compared to vehicle-treated animals (Fig. 6B and Suppl. Fig. 5).

We next evaluated H&E lung images to identify potential differences in the levels of SARS-CoV-2-induced syncytia between vehicle- and NTZ-treated hamsters. Strikingly, we observed multiple, often neighboring, syncytial epithelial cells in lungs from all the vehicle-treated SARS-CoV-2-infected animals at 5 dpi (Fig. 6C), which occurred exclusively in areas of bronchiole hyperplasia. By contrast, syncytia formation was rarely observed in the NTZ-treated group, and when observed consisted of a single isolated syncytium within an area of bronchiole hyperplasia (Fig. 6C). Notably, syncytia were not observed at either 2 or 14 dpi in any SARS-CoV-2-infected animals irrespective of treatment group; importantly, syncytia were never observed in naïve animals.

## DISCUSSION

A critical objective in the global effort to significantly reduce COVID-19 morbidity and mortality is the development of a scalable, well-tolerated, and cost-effective oral therapeutic or cocktail that can be deployed in resource-limited settings as well as in the global north. Here, we provide *in vitro* and *in vivo* data in physiological models of infection showing that NTZ significantly restricts the replication of different SARS-CoV-2 variants of concern *in vitro*, and inhibits virus-induced morbidity, inflammation, and viral dissemination *in vivo* in the Syrian hamster model. Together, these studies suggest that NTZ could have a potential role as an oral therapeutic in the treatment of COVID-19 and its long-term complications or as a prophylactic in high-risk exposures.

Multiple host processes have been implicated in the ability of NTZ to inhibit viral replication. In this respect, we previously showed that NTZ inhibition of EBOV replication is significantly impaired in cells deficient in the RNA sensors PKR and RIG-I (Jasenosky et al., 2019). EBOV viral protein 35 (VP35), a protein produced early in infection, binds to dsRNA replicative intermediates shielding EBOV from cytoplasmic sensors including RIG-I and PKR (Cardenas et al., 2006; Feng et al., 2007; Leung et al., 2009; Leung et al., 2010; Schumann et al., 2009), and a second EBOV protein, viral protein 24 (VP24), blocks the activation and nuclear accumulation of STAT1 and inhibits IFN-responsive gene transcription (Reid et al., 2006; Reid et al., 2007; Xu et al., 2014; Zhang et al., 2012). Thus, NTZ, at least in part, overrides EBOV mechanisms of viral innate immune evasion via its ability to amplify PKR- and RIG-I-dependent RNA sensing (Jasenosky et al., 2019). By contrast, the ability of NTZ to inhibit VSV infection is impaired in GADD34- and RIG-I-deficient cells, but not in PKR-deficient cells (Jasenosky et al., 2019). Thus, NTZ has the ability to inhibit viral replication by broadly amplifying RNA sensing pathways, but the specific host factors mediating its antiviral activity may differ during infection with different viruses (Jasenosky et al., 2019).

Here, we find that the antiviral activity of NTZ is enhanced by functional type-I IFN signaling. However, in the context of SARS-CoV-2 infection, NTZ was still able to significantly inhibit viral replication in type-I IFN-deficient Vero E6 cells and IFNAR-KO Ace2-A549 cells. It will be important in the future to assess whether the induction of type III IFN genes and the expression of a specific set of IFN-independent ISGs are contributing to this residual NTZ activity. Recent studies have shown that MDA5 and LGP2 are the major pattern recognition receptors involved in SARS-CoV-2 restriction (Sampaio et al., 2021; Yin et al., 2021). We previously showed that NTZ enhances the transcription and the signaling activity of MDA5 (Jasenosky et al., 2019; Ranjbar et al., 2019). However, the role of the MDA5-LGP2 sensing pathway in the antiviral activity of NTZ during SARS-CoV-2 infection remains to be defined and is currently a subject of investigation.

Interestingly, NTZ was previously shown to effectively inhibit the replication of influenza and parainfluenza viruses *in vitro* by impairing maturation and intracellular trafficking of their fusion glycoproteins (Piacentini et al., 2018; Rossignol et al., 2009). Furthermore, NTZ and niclosamide interact with TMEM family members TMEM16A and 16F, which are calcium-activated ion channels and lipid scramblases responsible for the exposure of phosphatidylserine on the cell surface (Braga et al., 2021; Caputo et al., 2008; Schroeder et al., 2008; Yang et al., 2008). Both drugs can function as TMEM16A antagonists, which attenuate bronchoconstriction (Miner et al., 2019). Moreover, niclosamide inhibits SARS-CoV-2 Spike-driven syncytia formation by antagonizing the activity of TMEM16F (Braga et al., 2021). Our results showing that NTZ broadly inhibits syncytia formation driven by SARS-CoV-2 Spike from major variants of concern are consistent with these studies and further broaden NTZ’s mechanism of action in viral infection. This finding is also consistent with the significant reduction of the number of SARS-CoV-2-infected cells in NTZ-treated cultures, and with the ability of NTZ to synergize with RDV given their independent mechanisms of action. It remains to be demonstrated whether or not NTZ alters spike maturation and intracellular trafficking, or if NTZ regulates the activity of members of the TMEM16 family in the context of infection.

Strikingly, oral administration of NTZ to SARS-CoV-2-infected Syrian hamsters results in a significant reduction of virus-induced weight loss and lung viral load. Consistently, NTZ treatment clearly affected Spike expression levels and distribution patterns within the lungs from infected hamsters. Furthermore, animals that had been treated with NTZ also exhibited a reduction of viral syncytial cells in their lungs. Whether this is directly due to the ability of NTZ to inhibit syncytia formation or is a consequence of the decreased viral load and spike expression is not yet known. Importantly, in addition to the significant decrease in spike dissemination, we also observed a dramatic decrease in bronchiolar epithelial degeneration/necrosis with reduced recruitment of inflammatory cells to bronchiolar lumina and lung interstitia at 2 dpi and to perivascular compartments at 14 dpi when hamsters who received NTZ recovered from infection. These results suggest that NTZ may inhibit injury to permissive cell types and ameliorate subsequent recruitment of inflammatory cells that could cause inflammatory complications of SARS-CoV-2 infection.

Intriguingly, an initial trial of NTZ treatment in patients (n=50) with moderate COVID-19 disease showed that 5 days of treatment was associated with decreased inflammatory biomarkers, faster SARS-CoV-2 PCR negativity, and decreased time to hospital discharge (Blum et al., 2021). Furthermore, a randomized controlled trial (RCT) (n=379) of 5 days of placebo or NTZ treatment for mild or moderate COVID-19 using the extended-release formulation of NTZ (NT-300) was associated with an 85% reduction in progression to severe illness (Rossignol et al., 2021). In both trials NTZ was well tolerated and there were no major adverse events. Here, our *in vivo* studies in the hamster model demonstrating that NTZ significantly reduces morbidity, inflammation, and viral dissemination are consistent with these studies.

Moreover, our demonstration that NTZ significantly inhibits SARS-CoV-2 growth, synergizes with RDV, and significantly inhibits different SARS-CoV-2 variants of concern including the more pathogenic delta (B.1617.2) variant, suggests that combining NTZ with another orally available antiviral could form a promising cocktail to treat SARS-CoV-2 infection. In this case, addition of NTZ to new oral antivirals shown to reduce hospitalization and death, such as molnupiravir or Paxlovid (Jayk Bernal et al., 2021; Ledford, 2021), or with intravenously delivered RDV, which has recently been shown to significantly decrease progression of disease in individuals at high risk (Gottlieb et al., 2021), could potentially mitigate viral spread in organs, inflammatory complications, and disease in high-risk exposures including immunosuppressed individuals where viremia is persistent and difficult to treat.

Finally, NTZ has a long safety profile in adults and children, has a pediatric syrup formulation, and is well-tolerated in HIV-infected individuals (Rossignol, 2014; Wang et al., 2020). An NTZ-based combination cocktail based on synergy could therefore potentially lower the dosage requirement of expensive and scarce new antiviral oral drugs that have a shorter history of safety data; this could be especially helpful in treatment of children and in immunosuppressed HIV-infected individuals. Furthermore, the oral formulation, inexpensive cost, and wide global use of NTZ make it easily deployable and accessible in resource-constrained settings.

## MATERIALS AND METHODS

### Cell lines

Vero E6 (ATCC, CRL-1586), A549 (ATCC, CRM-CCL-185)-derived, and HEK293T (ATCC, CRL-3216)-derived cell lines were maintained in DMEM (Corning) supplemented with 10% FBS (Peak Serum) and Penicillin/Streptomycin (Corning) at 37°C and 5% CO_2_. hACE2-A549 cells used in Fig. 2A (Daniloski et al., 2021) and hAce2-HEK293T (Miorin et al., 2020) were previously described. Vero TMPRSS2 (BPS Bioscience, 78081) cells were maintained in DMEM (Corning) supplemented with 10% FBS (Peak Serum), non-essential amino acids, sodium pyruvate (Corning), Penicillin/Streptomycin (Corning), and puromycin (3 μg/mL) (InvivoGen). IFNAR KO A549 cells were generated by CRISPR-Cas9 ribonucleoprotein (RNP) complex (IDT) transfection using the Nucleofector system (Lonza Bioscience). Specifically, a predesigned Alt-R CRISPR-Cas9 gRNA targeting exon 1 (design ID: Hs.Cas9.IFNAR1.1.AK), the ATTO 550 Alt-R CRISPR-Cas9 tracrRNA, and the Alt-R S.p. HiFi Cas9 Nuclease were used to form CRISPR-RNPs *in vitro*. Wild type and IFNAR KO A549 cells were then transduced with an Ace2 expression vector simultaneously using a pTRIP-SFFV-Blast-2A-myc-hACE2 construct, a kind gift from Nir Hacohen and Matteo Gentili (Massachusetts General Hospital and the Broad Institute), sequenced, and then functionally validated (Suppl. Fig. 3). All cell lines used in this study were regularly screened for mycoplasma contamination using the Universal Mycoplasma Detection Kit (ATCC, 30-1012K).

### Viruses

SARS-CoV-2 isolate USA-WA1/2020 (BEI Resources NR-52281) stocks were grown in Vero E6 cells as previously described (Miorin et al., 2020). hCoV-19/Japan/TY7-503/2021 (P.1) was obtained from BEI Resources (NR-54982). hCoV-19/USA/MD-HP01542/2021 JHU (B.1.351) was a kind gift from Dr. Andy Pekosz. The SARS-CoV-2 Delta variant (B.1.617.2 PV29995) was obtained from Dr. Viviana Simon (Mount Sinai Pathogen Surveillance Program). B.1.351, B.1.617.2, and P.1 viral stocks were grown on Vero TMPRSS2 cells. All viruses were validated by genome sequencing. NG-SARS-CoV-2, a stable monomeric NeonGreen-labeled SARS-CoV-2 virus (icSARS-CoV-2-mNG), was a kind gift from Pei-Yong Shi (University of Texas Medical Branch) (Xie et al., 2020). Virus was reconstituted from lyophilized sample and then titered using Vero cells as described (Huang et al., 2020; Xie et al., 2020). All experiments performed with a SARS-CoV-2 isolate were performed in accordance with biosafety protocols developed by ISMMS, the Ragon Institute of MGH, MIT and Harvard, or the National Emerging Infectious Diseases Laboratories of Boston University.

### *In vitro* antiviral growth assay

Vero E6, Ace2-A549, or Ace2-HEK293T cells were seeded in 96-well plates in DMEM (10% FBS) and incubated for 24 h at 37°C and 5% CO_2_. Four hours before infection the medium was replaced with 100 μl of DMEM (2% FBS) containing the compound of interest (NTZ, TIZ, or remdesivir) at concentrations 50% greater than those indicated, including a DMSO control. Plates were then transferred into the BSL-3 facility and infected at the desired MOI in 50 μl of DMEM (2% FBS), bringing the final compound concentration to those indicated. Plates were then incubated for 48 h at 37°C. After infection, supernatants were removed, and cells were fixed with 4% paraformaldehyde for 24 hours prior to being removed from the BSL-3 facility. The cells were then immunostained for the viral NP protein (an in-house mAb 1C7 provided by Dr. Thomas Moran, Mount Sinai School of Medicine) with a DAPI counterstain. Infected cells (488 nm) and total cells (DAPI) were quantified using a Celigo (Nexcelcom) imaging cytometer. Percent infection was quantified as ([infected cells/total cells]-background) multiplied by 100 and the DMSO control was set to 100% infection for analysis. The IC_50_ was determined using GraphPad Prism software. Cytotoxicity was performed using the MTT assay (Roche) in uninfected cells with the same compound dilutions and concurrent with the viral replication assay. All assays were performed in biologically independent triplicates. Remdesivir was purchased from Medkoo Biosciences, Inc.

### *In vitro* viral growth and infection analysis by FACS and Imaging Flow Cytometry

For standard flow-cytometric analyses, cells treated with NTZ or mock treated and infected with NG-SARS-CoV2 and were terminated at the indicated time points. Cells were washed with PBS, fixed with 4% paraformaldehyde for 24 hrs and acquired on a BD FACSCanto. NeonGreen fluorescent infected cells were evaluated using Flow-Jo software after gating on live cell populations. For ImageStream analysis, cells treated with NTZ or mock treated and infected with NG-SARS-CoV2 were terminated at the indicated time points. Cells were washed with PBS and fixed with 4% paraformaldehyde for 24 hrs. Next, 3000 gated cells were acquired on an Amnis ImageStream^X^ Mark II imaging flow cytometer (EMD Millipore). Live focused single cells were gated and NeonGreen fluorescent positive cells were evaluated using Ideas software and pixels were exported for further analysis on Prism to graph the dot plots and compare NTZ-treated to mock by the Mann-Whitney two-tailed test. To compare the numbers of NTZ-treated infected cells to mock-treated cells we used the Fisher exact test using Stata 12 software.

### H9 stem cell-derived pneumocyte-like cell differentiation and SARS-CoV-2 infection

Human embryonic stem cells (H9, WiCell, WA09) were cultured with mTeSR (STEMCELL Technologies, 85850) on Vitronectin XF (STEMCELL Technologies, 07180)-coated tissue culture plates and split in a ratio of 1:6 to 1:12 every four to six days with Gentle Cell Dissociation Reagent (STEMCELL Technologies, 07174). Alveolar differentiation was induced as previously described (Riva et al., 2020; White et al., 2021). Viral infections were performed in the BSL-3 facility on day nine after induction of differentiation, in accordance with biosafety protocols developed by ISMMS. Cells were fixed and analyzed two days post-infection. For assessment of infection rates, H9 cells were washed once with PBS, detached with 5 mM EDTA for 5 min at 37°C, and then fixed with 4% paraformaldehyde. Cells were washed in PBS supplemented with 2 mM EDTA, permeabilized using Perm/Wash buffer (BD Biosciences), and subsequently stained for 1 h at room temperature with anti-NP mAb 1C7 antibody labeled with an Alexa488-fluorescent marker using the Alexa Fluor™ 488 Antibody Labeling Kit (Invitrogen). Immunofluorescence-labelled cells were washed twice in PBS-EDTA after incubation with the antibody and then subjected to flow-cytometry analysis using a Gallios flow cytometer (Beckman Coulter). Cytotoxicity of the compounds was assessed using the MTT assay (Roche) in uninfected cells treated with the same compound dilution and concurrent with the viral replication assay.

### iPSC differentiation into alveolar epithelial type 2 cells (iAT2s) for air-liquid interface culture and SARS-CoV-2 infection

Previously published human iPSCs (Huang et al., 2020) carrying an SPC2-SFTPC^tdTomato^ reporter (iPSC clone SPC2-ST-B2, Boston University (Hurley et al., 2020)) were differentiated into alveolar epithelial type 2 cells (iAT2s) and maintained as alveolospheres embedded in three-dimensional (3D) Matrigel in “CK+DCI” media, as previously described (Jacob et al., 2019). iAT2s were serially passaged approximately every 2 weeks by dissociation into single cells via the sequential application of dispase (2 mg/mL; Thermo Fisher Scientific; 17105-04) and 0.05% trypsin (Invitrogen; 25300054) and replated at a density of 400 cells/μL of Matrigel (Corning; 356231), as previously described (Jacob et al., 2019). To generate air-liquid interface (ALI) cultures, iAT2s were seeded at 520,000 live cells/cm^2^ on Transwell inserts (Corning Cat#3470), and apical media was removed 2 days post-seeding. After 7 days post-seeding, iAT2 ALI cultures were ready to be infected with SARS-CoV-2 USA_WA1/2020 or SARS-CoV-2-mNG at an MOI of 0.04, along with mock controls. Prior to infection, RDV (Selleck Chem Cat#S8932) or NTZ was added to the apical and basolateral sides of the Transwell inserts for 30 minutes as indicated. The apical side was subsequently aspirated and replaced by SARS-CoV-2 diluted in CK+DCI media for one hour. At 2 days post-infection, samples were collected for downstream processing.

For flow cytometry and trypan blue staining, 200 μL of Accutase (Sigma Cat# A6964) was added apically and incubated for 20 minutes at room temperature. The cells were collected and used directly for trypan blue staining or fixed with 10% formalin for at least 6 hours at 4°C and subsequently removed from BSL-4 prior to FACS analysis. After single cell suspension, analysis was performed by immunostaining for the viral NP protein as described for the H9-pneumocytes or by imaging flow cytometry using the ImageStream platform. For fluorescence microscopy, iAT2 ALI cultures were fixed with 10% formalin for at least 6 hours at 4°C at the indicated times post-infection. Samples were then removed from BSL-4 and washed with PBS three times for 5 minutes at room temperature. Cell nuclei on the transwell inserts were stained with DAPI dye (1:500, Life Technologies). The Transwell insert membranes were then cut out and mounted on slides using ProLong Diamond Antifade Mountant (Invitrogen Cat#P36965). Slides were imaged on a confocal microscope (Leica SP5).

### Antiviral combination assay

2,000 Vero E6 cells per well were seeded into 96-well plates in DMEM (10% FBS) and incubated for 24 h at 37°C and 5% CO_2_. Two hours before infection, the medium was replaced with 100 μl of DMEM (2% FBS) containing the combination of NTZ and RDV following a dilution combination matrix. A 6 by 6 matrix of drug combinations was prepared in triplicate by making serial two-fold dilutions of the drugs on each axis, including a DMSO control column and row. The resulting matrix had no drug in the right upper well, a single drug in rising 2-fold concentrations in the vertical and horizontal axes starting from that well, and the remaining wells with rising concentrations of drug mixtures reaching maximum concentrations of the drugs at the lower left well. Plates were then transferred into the BSL-3 facility and SARS-CoV-2 (MOI 0.025) was added in 50 μl of DMEM (2% FBS), bringing the final compound concentration to those indicated in the figures. Plates were then incubated for 48 h at 37ºC. After infection, cells were fixed with formaldehyde for 24 hours prior to being removed from the BSL-3 facility. The cells were then immunostained for the viral NP protein as described above. This combination data was analyzed using SynergyFinder (Ianevski et al., 2020) by the Loewe method (Loewe, 1953).

### Spike-induced syncytia assay

30,000 Vero-TMPRSS2 cells were reverse transfected with 100 ng of pCAGGS-S plasmids in a black 96-wellplate using TransIT-LT1 Transfection reagent (Mirus) in complete growth medium (containing 10% FBS). 4 hrs after transfection, medium containing transfection complexes was removed and replaced by medium containing 2% FBS and either DMSO or 15 μM NTZ. Cells were incubated for 24 hrs at 37°C to allow syncytium formation. After 24 h, cells were fixed with 4% paraformaldehyde in PBS (15 min), permeabilized with 0.1% Triton X-100 (15 min) and stained for 30 min with 2 μg/ml HCS Cell Mask stain (Life Technologies). Syncytia were counted microscopically in triplicate wells. The pCAGGS-S plasmids used in this assay have been previously described (Escalera et al., 2021).

### *In vivo* Syrian hamster infection with SARS-CoV-2

Syrian Golden hamsters 10-12 weeks old (Envigo RMS, LLC), n = 42, were divided into 3 groups of 14: naïve (n = 14), SARS-CoV-2-infected mock (PBS) treatment cohort (n = 14), and SARS-CoV-2-infected NTZ (NT-300, a gift from Romark, Inc)-treated at 300 mg/kg/per day bi-daily cohort (n = 14). Hamsters in the infected cohorts were infected intranasally with SARS-CoV-2 (isolate USA-WA1/2020) in a 50 μl suspension containing a targeted dose of 100 PFU (actual dose was 300 PFU). On the day of SARS-CoV-2 challenge the mock cohort was treated with sterile PBS and the NTZ-treated cohort was treated with 150 mg/kg of NTZ at 6 hours before challenge and then 6 hours after challenge with subsequent treatments for these 2 cohorts every 12 hours (bi-daily) for the remaining 4 days of treatment. Cohorts were treated via oral gavage using an 18 gauge x 2 inch curved animal feeding needle to deliver 1 ml of volume per treatment. Lungs and sera were harvested for virus load analysis from 4 hamsters per group on day 2, 5, and 14 post SARS-CoV-2 challenge. Weight loss was monitored from the first treatment time point through end of the study on day 14 post SARS-CoV-2 challenge. Weight loss was monitored for all hamsters throughout the study for humane endpoint criteria. Percent weight loss data (Fig. 4A) were derived from 6 hamsters that went to study endpoint at day 14 post-SARS-CoV-2 challenge.

### Plaque assay

Plaque assays were performed using Vero E6 cells as previously described (Amanat et al., 2020). Briefly, Vero E6 cells seeded in 12-well plate format were infected with serial ten-fold dilutions of supernatants from homogenized lung tissues. Virus absorption was carried out for 1 hour using an inoculum of 200 μl and rocking the plates every 10-15 min. After 1 hour, the inoculum was removed, and the cells were incubated with an overlay composed of MEM with 2% FBS and 0.7% Oxoid agar for 72 hours at 37°C with 5% CO_2_. The plates were subsequently fixed using 5% formaldehyde and immuno-stained using a monoclonal anti-SARS-CoV-NP antibody (Creative-Biolabs; NP1C7C7). In brief, plates were blocked (3% skim-milk TBS with 0.1% Tween20 for 1 h), stained for 90 min with anti-NP antibody (mAb 1C7, diluted 1:1000 in 1% skim-milk TBS with 0.1% Tween20), and finally secondary-stained with anti-mouse-HRP (antibody diluted 1:5000 in 1% skim-milk TBS with 0.1% Tween20 for 45 mins). Plates were incubated for 10 min with KPL TrueBlue peroxidase substrate (Seracare) to reveal staining.

### Pathology analysis: histologic processing and semi-quantitative analysis

Tissue samples were fixed for a minimum of 7 days in 10% neutralized buffered formalin before being removed from BSL-4 and subsequently processed in a Tissue-Tek VIP-6 automated vacuum infiltration processor (Sakura Finetek) and embedded in paraffin using a HistoCore Arcadia paraffin embedding machine (Leica). 5 μm tissue sections were generated using an RM2255 rotary microtome (Leica) and transferred to positively charged slides, deparaffinized in xylene, and dehydrated in graded ethanol. Tissue sections were stained with hematoxylin and eosin for histologic examination, with additional serial sections utilized for immunohistochemistry (IHC). A Ventana Discovery Ultra (Roche) tissue autostainer was used for IHC. Specific protocol details are outlined in Suppl. Table 1. In brief, SARS-CoV-2 spike protein monoplex IHC was conducted using a Chromomap DAB IHC kit (Roche, Basel, Switzerland) with CC1 antigen retrieval at 95°C for 64 minutes, primary incubation at 1:900 for 40 min at room temperature, rabbit anti-IgG1+IgG2a+IgG3 antibody (ab133469) at 37°C for 20 minutes (1:1,000), and ImmPRESS HRP Goat Anti-Rabbit IgG polymer pre-dilute detection (Vector Laboratories) at 37°C for 20 minutes. Uninfected animals served as negative controls, while SARS-CoV-2 animals receiving PBS and no therapeutic served as positive controls confirming the specificity of the assay. Histomorphological analysis was performed by a single board-certified veterinary pathologist (N.A.C.), who developed an ordinal grading score encompassing the diversity and severity of histologic findings outlined in Table 2. Histologic criteria were broken down into three compartments: airways, blood vessels, and interstitium, with results utilized to generate a cumulative lung injury score. This score also incorporated the overall degree of immunoreactivity to the SARS-CoV-2 Spike antigen. A summary of individual animal scores and specific criteria utilized to score lungs is included in Suppl. Tables 1 and 2.

### Quantitative Image Analysis

Digitized whole slide scans were analyzed using the image analysis software HALO (Indica Labs, Inc., Corrales, NM). Slides were manually annotated to select pulmonary parenchyma, excluding non-pulmonary tissues (i.e., tracheobronchial lymph nodes, adipose tissue, etc.). Quantitative outputs were obtained using the Area Quantification (AQ) module, which reports total area of immunoreactivity of a specified parameter within a region of interest. Values are given as a percentage of total tissue area analyzed. Minimum dye intensity thresholds were established using the real-time tuning field of view module to accurately detect positive immunoreactivity for SARS-CoV-2 Spike antigen.

## ACKNOWLEDGEMENTS

This work was funded by grants from the Annenberg Foundation, Fast Grants, and the NIH (R21AI151732), and a laboratory gift from Jeanne Sullivan to AEG. This research was also partly funded by grants from CRIPT (Center for Research on Influenza Pathogenesis and Transmission), an NIAID-funded Center of Excellence for Influenza Research and Response (CEIRR, contract 75N93021C00014), NCI SeroNet grant U54CA260560; NIAID grants U19AI135972 and U19AI142733, DARPA grant HR0011-19-2-0020; a supplement to DoD grant W81XWH-20-1-0270, JPB Foundation, Open Philanthropy Project (research grant 2020-215611 (5384)), and by anonymous donors to AG-S; and by PRIME (Program for Research on Immune Modeling and Experimentation), an NIAID-funded Modeling Immunity for Biodefense Center (grant U19 AI117873), and CRIP (Center for Research on Influenza Pathogenesis), and an NIAID-funded Center of Excellence for Influenza Research and Surveillance (CEIRS, contract HHSN272201400008C) to AF-S and AG-S. The work was also supported by grants from the NIH (R01HL095993, U01TR001810, N01 75N92020C00005) and Evergrande COVID-19 Response Fund Awards from the Massachusetts Consortium on Pathogen Readiness (MassCPR) to DNK; Fast Grants, NIH (NCATS, UL1TR001430), and MassCPR to EM; MassCPR to RC; SIG grants S10-OD026983 and S10-OD030269 to NAC; NIH (F30HL147426) to KMA; CJ Martin Early Career Fellowship from the Australian National Health and Medical Research Council to RBW; and NIH (U01TR001810, UL1TR001430, R01DK101501, and R01DK117940) to AAW.

We thank Romark Inc. for the gift of NTZ-300, S-Y Pei and UTMB for the gift of NG-SARS-CoV-2, Nir Hacohen and Matteo Gentili for the gift of pTRIP-SFFV-Blast-2A-myc-hACE2, and Michael Farzan and Huihui Mu for the gift of pQCXIP-myc-hACE2-c9. We also thank Randy Albrecht for support with the BSL-3 facility and procedures at the ISMMS, Julie Boucou and Xie Yong for support and advice at the BSL-3 facility at the Ragon Institute of MGH, MIT and Harvard, Richard Cadagan, Hans P. Gertje, and Paige Montanaro for technical assistance, and the Microscopy Shared Resource Facility at the Icahn School of Medicine at Mount Sinai. We are grateful to Gail Cassell for helpful discussions through the years. Finally, we are indebted to Wallis Annenberg, Jeanne Sullivan, and the founders of Fast Grants for their critical and early support.

