## Supplementary Materials for "The oral drug nitazoxanide restricts SARS-CoV-2 infection and attenuates disease pathogenesis in Syrian hamsters"

### SUPPLEMENTARY FIGURE LEGENDS

**SFig 1: Confocal microscopy images of Vero E6 (A) or Ace2-A549 (B) cells infected with SARS-CoV-2.** At 24 hrs post-infection, cells were stained with antibodies against NP or against the dsRNA intermediate of replication (Scale bars, 20 µm).

**SFig 2. SARS-CoV-2 infects Ace2-HEK293T cells and replication is inhibited by NTZ.** Concentration-response curves for viral infectivity and for cell viability in Ace2-HEK293T cells treated with NTZ or RDV as described in the Figure 1 legend. Cells were infected with SARS-CoV-2 (isolate USA-WA1/2020) at an MOI of 0.25.

**SFig 3. Functional validation of Ace2-A549 IFNAR-KO cells.**

**A.** WT and IFNAR-KO Ace2-A549 cells were treated for 45 minutes with 1,000 U/ml of type I IFN before lysis. Expression of the indicated protein was determined by Western blot using actin as a loading control. **B.** WT and IFNAR-KO Ace2-A549 cells were stimulated for 24 hrs with 100 U/ml of type I IFN. The expression of IFITM3 mRNA was determined by qRT-PCR. IFITM3 mRNA levels relative to cyclophilin are shown.

**SFig 4. Infection of IAT2 cells with NG-SARS-CoV-2 shown by confocal microscopy.** At 7 days post-plating 400,000 iAT2s/transwell were infected with NeonGreen-SARS-CoV-2 (NG-SARS-CoV-2) at an MOI of 0.01 for 2 days, fixed, and evaluated by confocal microscopy.

**SFig 5. High magnification of images shown in Figure 6 at day 2 and 14 of lung from naïve and from SARS-CoV-2+PBS/vehicle- vs. SARS-CoV-2+NTZ-treated animals at day 2 and 14 post-infection. Top: Bronchioles.** At 2 dpi SARS-CoV-2 PBS/vehicle-treated: bronchioles (black arrows) are occluded by necrotic cellular debris and neutrophils and histiocytes, with neighboring bronchiole epithelial apoptosis and/or necrosis represented by nuclear fragmentation (black hashed boxes). At 2 dpi SARS-CoV-2 NTZ-treated: segmental bronchiole epithelial degeneration and denuding represented by a hashed box, with less overall luminal exudate. **Bottom: Interstitium/Blood Vessels.** At 14 dpi SARS-CoV-2 PBS/vehicle-treated: mild-to-moderate residual perivascular lymphocytic infiltrate (see black box inset) neighboring areas of residual Alveolar type 2 (AT2) cell hyperplasia (blue arrows). At 14dpi SARS-CoV-2 NTZ-treated: minimal sporadic residual perivascular lymphocytic infiltrate (see black box inset at higher magnification) neighboring areas of residual AT2 cell hyperplasia (blue arrows). Scale bars: top row (bronchioles), 50 μm; bottom row: interstitium/blood vessels, 100 μm.

**S Table 1. Monoplex SARS-CoV-2 Spike DAB Immunohistochemistry (IHC)**

**S Table 2. Lung Ordinal Scoring System**

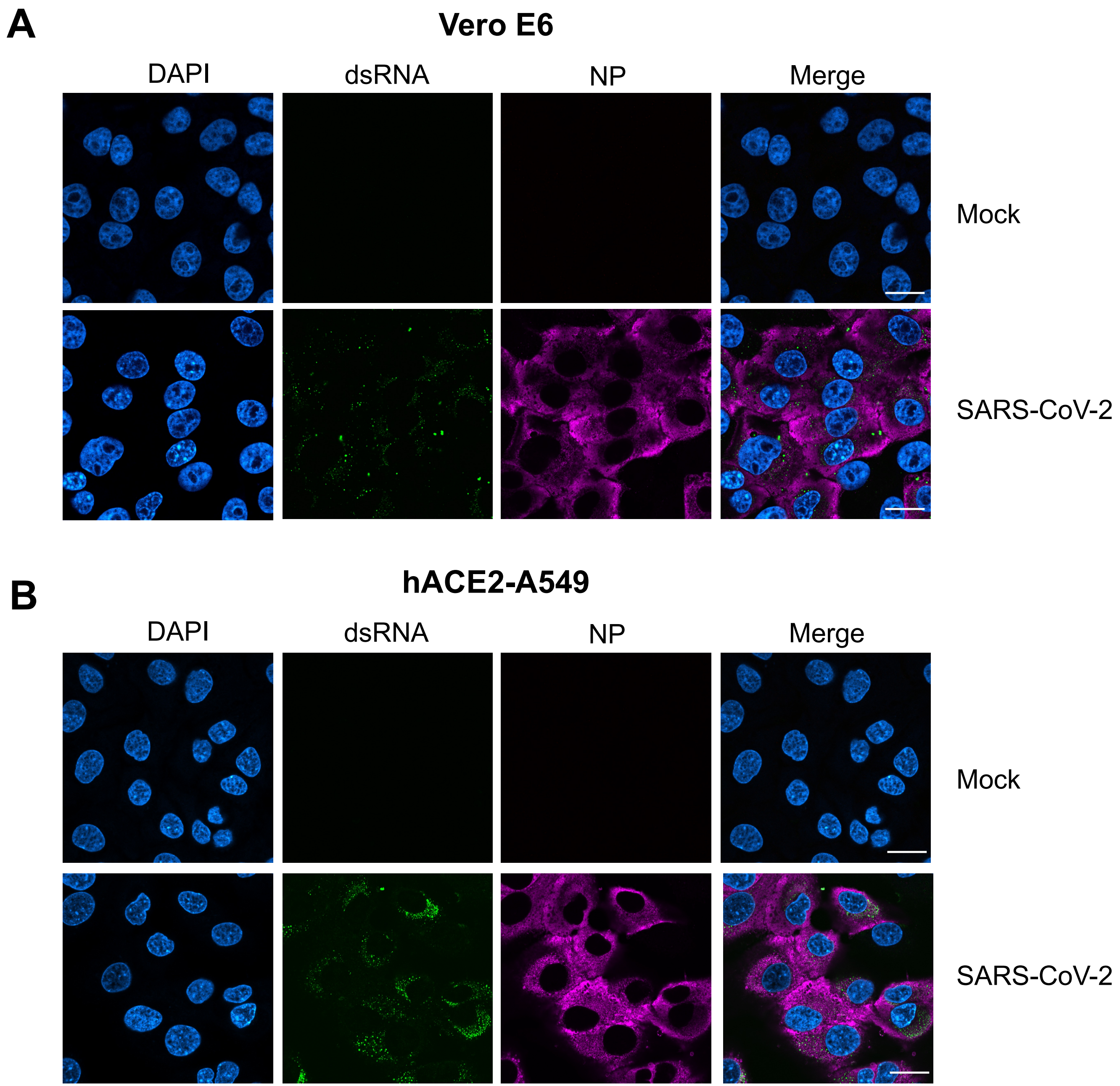
A
Vero E6
DAPI
dsRNA
NP
Merge
Mock
SARS-CoV-2
B
hACE2-A549
DAPI
dsRNA
NP
Merge
Mock
SARS-CoV-2

S. Fig. 1

**Ace2-HEK293T / SARS-CoV-2-WA1/2020**

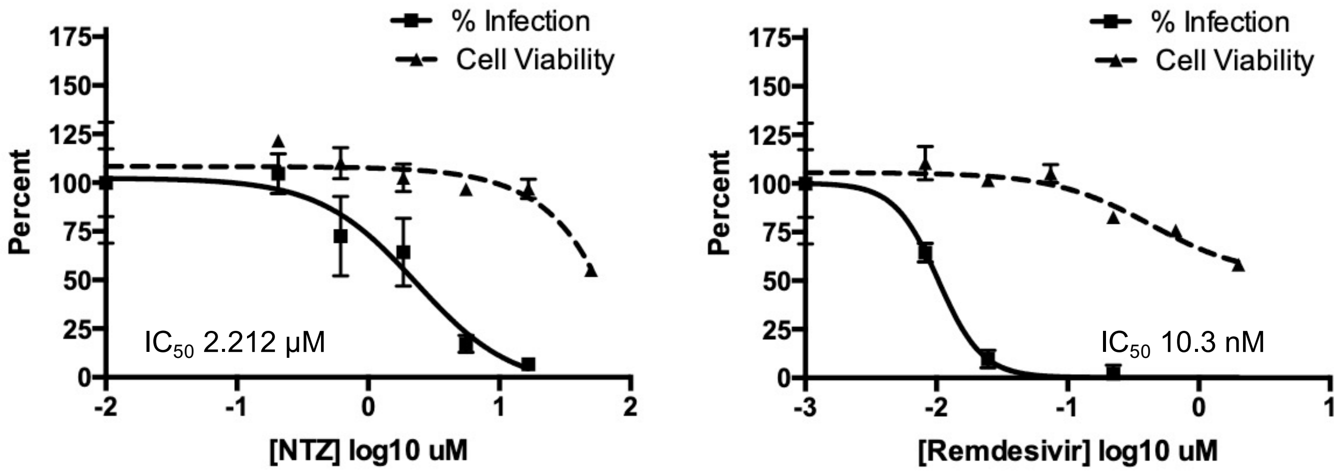


S. Fig. 2

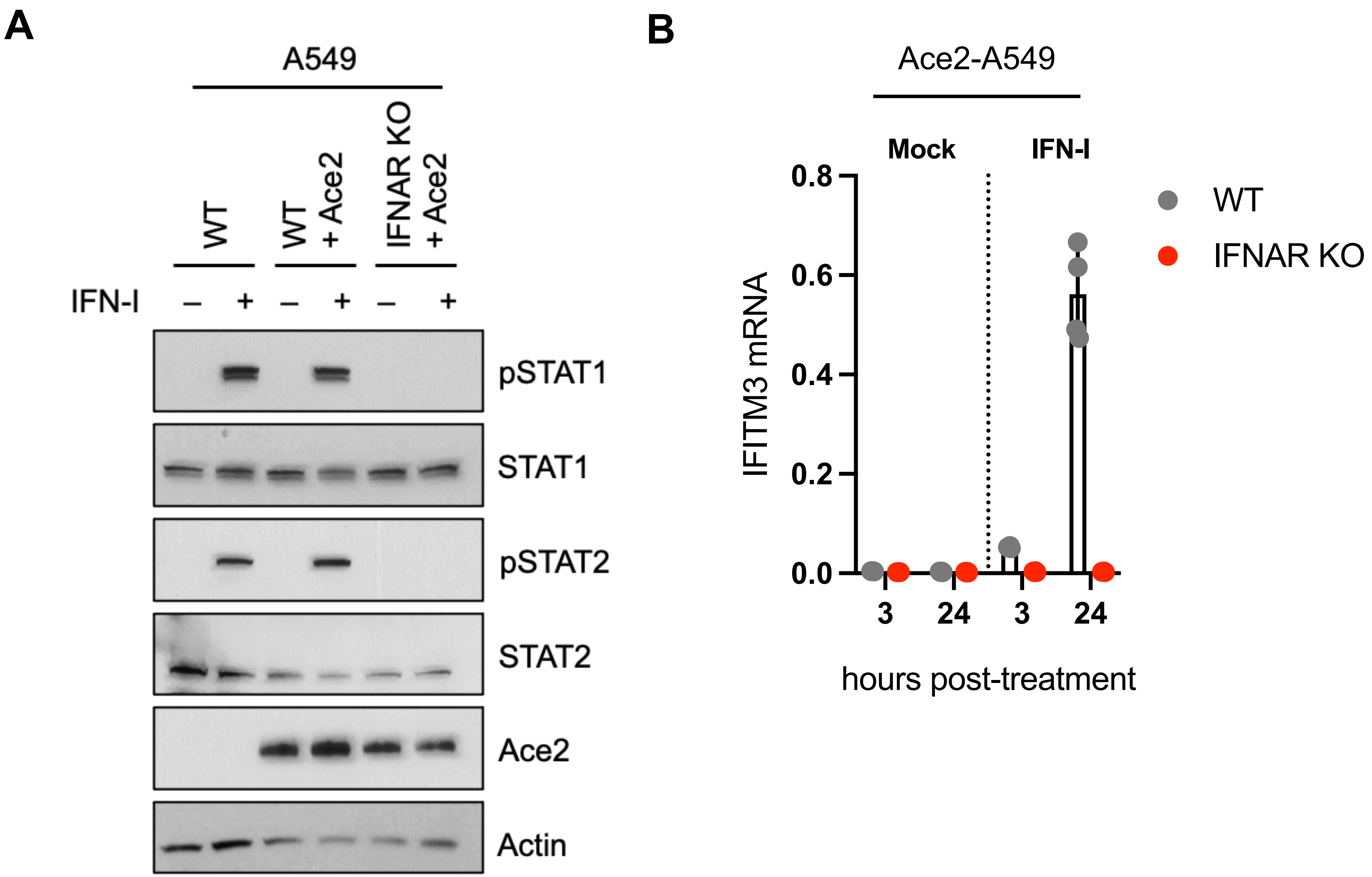
A
A549
WT
WT + Ace2
IFNAR KO + Ace2
IFN-I – + – + – +
pSTAT1
STAT1
pSTAT2
STAT2
Ace2
Actin
B
Ace2-A549
Mock
IFN-I
IFITM3 mRNA
0.8
0.6
0.4
0.2
0.0
3
24
3
24
hours post-treatment
WT
IFNAR KO

S. Fig. 3

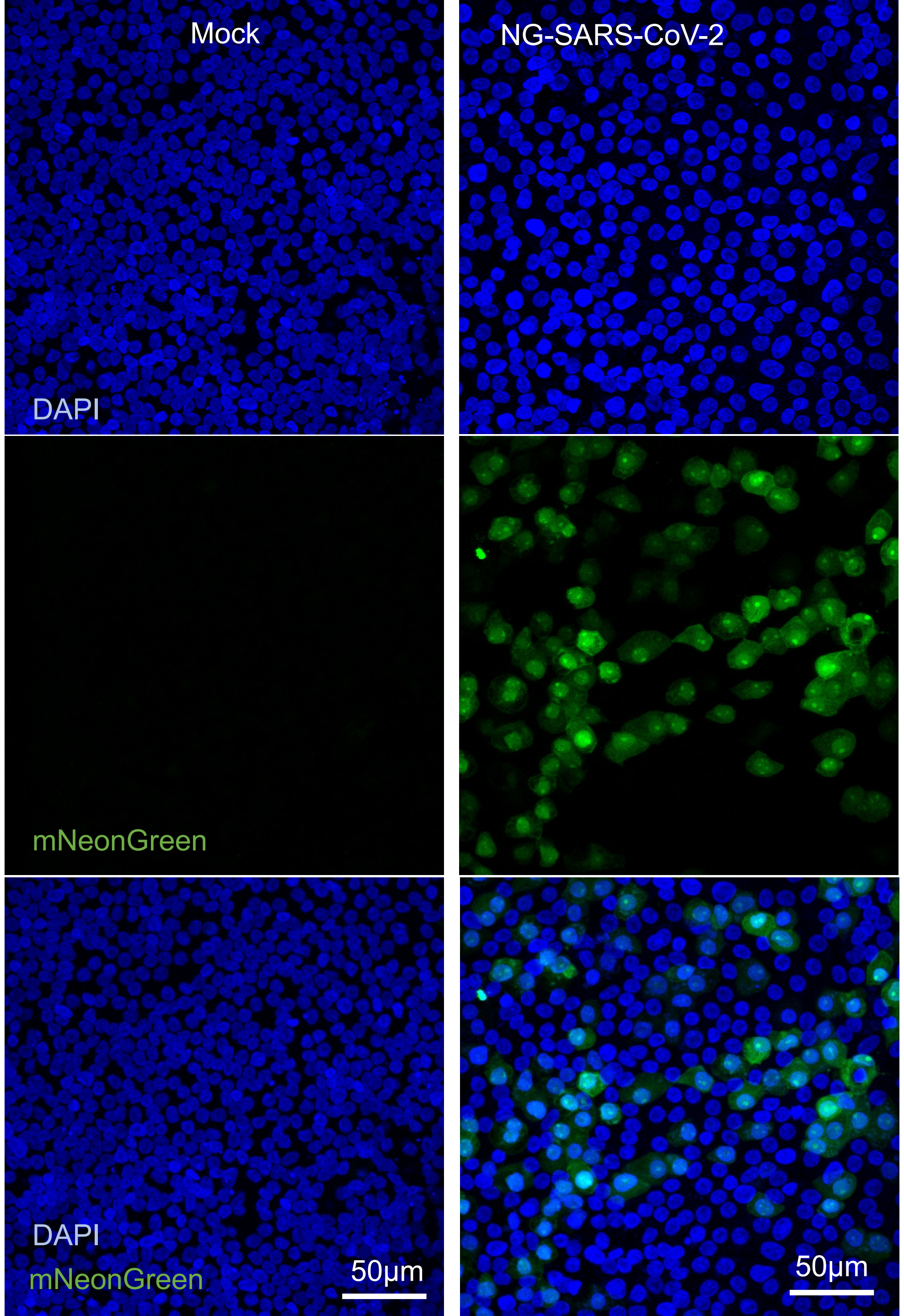

Mock
NG-SARS-CoV-2
DAPI
mNeonGreen
DAPI
mNeonGreen
50μm
50μm


S. Fig. 4

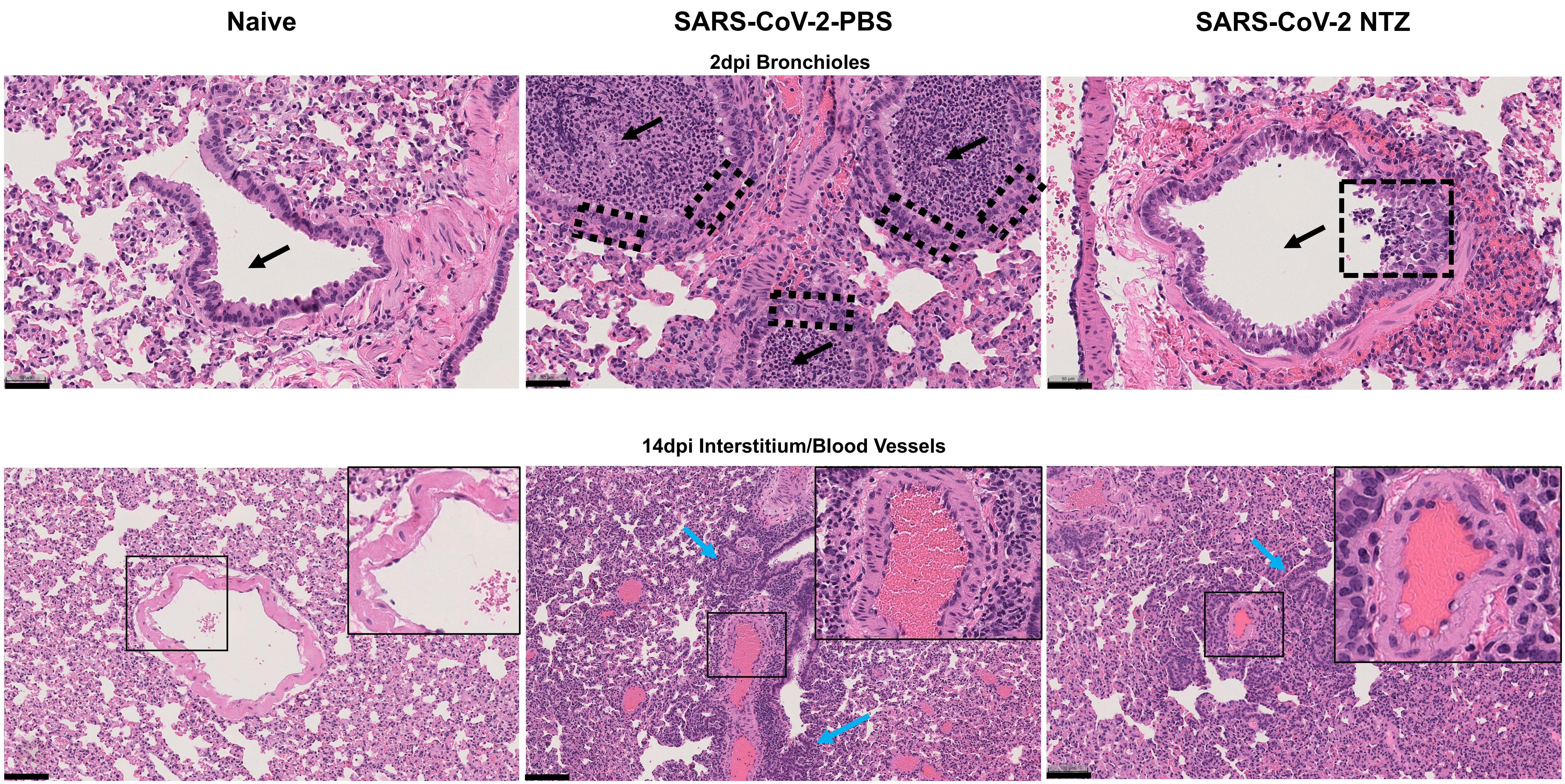
Naive
SARS-CoV-2-PBS
SARS-CoV-2 NTZ
2dpi Bronchioles
14dpi Interstitium/Blood Vessels

S. Fig. 5

### Supplementary Tables

| S Table 1. Monoplex SARS-CoV-2 Spike DAB Immunohistochemistry (IHC) | | | | |
|---|---|---|---|---|
| Species | Antigen | Manufacturer and Clone | Primary Antibody Dilution | Chromogen |
| Ms monoclonal | SARS-CoV-2 Spike | Cell Signaling Technology E7U60 | 1:900 | DAB |
| Diaminobenzidine-DAB; IHC-immunohistochemistry | | | | |

| **Suppl. Table 2. Lung Ordinal Scoring System** | |
|---|---|
| **Airways (bronchioles)** | |
| 0 | Within normal limits. |
| 1 | Mild bronchiole epithelial degeneration/necrosis and hyperplasia/hypertrophy; mononuclear +/- neutrophil peribronchiolar and/or airway infiltrates |
| 2 | Moderate bronchiole epithelial degeneration/necrosis and hyperplasia/hypertrophy; mononuclear +/- neutrophil peribronchiolar infiltrates; infiltration of bronchiole airways with moderate mononuclear infiltrates, neutrophils, necrotic cellular debris, +/- erythrocytes. |
| 3 | Marked to severe bronchiole epithelial degeneration/necrosis and/or hyperplasia/hypertrophy with syncytial cells; mononuclear +/- neutrophil peribronchiolar infiltrates; +/- infiltration of bronchiole airways with mononuclear infiltrates, neutrophils, necrotic cellular debris, +/- erythrocytes. |
| Interstitium | |
| 0 | Within normal limits. |
| 1-A (acute) | Mild focal to multifocal interstitial expansion and/or alveolar infiltration by mononuclear cells +/- neutrophils, with no observable edema, hemorrhage, and/or fibrin exudation. |
| 1-C (chronic) | Mild to moderate alveolar type 2 (AT2) pneumocyte hyperplasia, with mild interstitial or alveolar mononuclear infiltrates, +/- pleural and/or interstitial fibrosis. |
| 2-A (acute) | Moderate multifocal interstitial expansion and/or alveolar infiltration by mononuclear cells (histiocytes and/or lymphocytes) and neutrophils, with mild edema, hemorrhage, and/or fibrin exudation, admixed with low amounts of cellular debris. |
| 2-C (chronic) | Mild to moderate multifocal interstitial expansion and/or alveolar infiltration by mononuclear cells +/- neutrophils, with mild to moderate AT2 pneumocyte hyperplasia |
| 3-A (acute) | Marked to severe multifocal interstitial expansion and/or alveolar infiltration by mononuclear cells and neutrophils, with moderate to regionally severe edema, hemorrhage, and/or fibrin exudation, with mild to moderate cellular debris. |
| 3-C (chronic) | Severe alveolar type 2 (AT2) pneumocytes hyperplasia +/- alveolar bronchiolization. Prominent anisocytosis and cytomegaly of AT2 pneumocytes, with increased mitotic figures. Moderate to marked multifocal interstitial expansion and/or alveolar infiltration of mononuclear (histiocytes and/or lymphocytes) and neutrophils with mild edema, hemorrhage, and/or fibrin exudation, and mild cellular debris. |
| Blood vessels | |
| 0 | Within normal limits. |
| 1 | Mild mononuclear perivascular infiltrate +/- reactive endothelial hypertrophy. |
| 2 | Moderate mononuclear perivascular infiltrate +/- neutrophils, prominent reactive endothelium, +/- margination and/or transmigration of leukocytes. |
| 3 | Marked mononuclear infiltrate +/- neutrophils, prominent perivascular edema, reactive endothelium, and +/- margination and/or transmigration of leukocytes. |
